# The modular evolution of chitinases is governed by coevolution of auxiliary and catalytic domains

**DOI:** 10.64898/2026.01.15.699799

**Authors:** S. B. Pulsford, J. A. T. Dalton, O. B. Smith, J. Horton, R. L. Frkic, J. Z. Chen, J. Kaczmarski, C. J. Jackson

**Author notes:** Correspondence to: Pulsford SB or Jackson CJ. Joint first-author (S. B. Pulsford, J. A. T. Dalton).

## Abstract

The modular architecture of multi-domain enzymes is a key source of functional diversity. GH18 chitinases exemplify this, possessing a variety of auxiliary domains that have been postulated to underpin their role as essential global nutrient cyclers. However, there is no evolutionary framework that accounts for vast the assortment of domain compositions observed across the protein family. Here, we map the sequence space of nearly 40,000 bacterial chitinases, integrating phylogenetics with a quantitative analysis of domain promiscuity to decode their evolutionary history. We reveal that diversification follows distinct trajectories: enzymes necessary for environmental chitin scavenging evolve via high modular plasticity and structural elaboration of the catalytic core, whereas sequences required for specialized, essential developmental roles fix their domain architecture to fulfill lineage-specific physiological roles. By resolving how domain accretion co-evolves with catalytic scaffolds, this work provides a framework for understanding how multi-domain enzymes adapt to complex environments.

## Introduction

Chitin is a fundamental structural component in fungal cell walls and arthropod exoskeletons and is the second most abundant polysaccharide in nature, representing an enormous reservoir of renewable biomass^1^. Its enzymatic breakdown is primarily mediated by chitinases, which hydrolyze β-1,4-glycosidic bonds between N-acetylglucosamine (GlcNAc) monomers to produce oligomeric, dimeric, and monomeric breakdown products^2^. Bacterial chitinases play pivotal roles in global biogeochemical cycles, with chitinolytic bacteria in marine ecosystems processing significant portions of chitin turnover, while soil microbial communities rely on chitinase activity to release nitrogen and carbon from fungal biomass, underpinning terrestrial nutrient cycling^3,4^. Beyond their ecological significance, many of their breakdown products have valuable applications^1,5^. As demand for chitin-derived products continues to grow, there is an unmet need to discover and engineer chitinases with enhanced specificity and catalytic efficiency to overcome these limitations.

The CAZy (Carbohydrate Active Enzymes) database categorizes chitinases into three glycoside hydrolase (GH) families, GH18, GH19, and GH20, based on the sequence and structural homology of their primary catalytic domain^6,7^. Among these, the GH18 family is the largest and most enzymatically diverse family^6,8^. These enzymes degrade chitin while retaining the configuration of the β-anomeric carbon of substrates through a substrate-assisted double displacement hydrolytic mechanism^9,10^. The conserved DxxDxDxE catalytic motif, of which the glutamate residue acts as a general acid/base, is involved in protonating the glycosidic oxygen of the sugar substrate, which initiates the hydrolysis of the glycosidic bond^11^. While GH18 enzymes broadly share this mechanism, they display significant variation in substrate specificity; for instance, they may have varying degrees of exo-activity where they cleave substrates from one end of the polymer, and/or endo-activity in which random cleavage of glycosidic bonds occurs along the chitin chain^12,13^. Another key feature is processivity, i.e., the ability to catalyse multiple successive reactions while attached to the substrate; processive chitinases are characterized by deep clefts lined with conserved aromatic residues that bind individual chains and successively cleave off sugar monomers or dimers^14–16^. In contrast, non-processive variants detach after each hydrolytic reaction^17^. This functional variation allows chitin-degrading organisms to secrete a synergistic cocktail of GH18 proteins for cooperative breakdown of complex polymers^14^.

GH18 enzymes feature certain structural insertions and auxiliary domains that affect their activity^18^. The chitinase insertion domain (CID) is an insertion between the seventh ɑ-helix and seventh β-strand of the TIM barrel catalytic domain, and is proposed to enhance substrate association and potentially exo-chitinase activity by creating a deeper binding cleft^19^. Similarly, auxiliary domains are hypothesized to influence substrate affinity and catalytic efficiency^20,21^. Removal of these domains typically results in active enzymes with impaired binding to polymeric substrates^22^. Moreover, many organisms are observed to express multiple chitinases with diverse structural motifs and auxiliary domain composition, supporting the notion these structural variants have a broad spectrum of complementary functional properties^23^. For example, the soil bacterium *Streptomyces coelicolor* encodes over 10 chitinase genes with diverse architectures^24^. This is consistent with these prominent structural/sequence features acting as major contributors to GH18 functional diversification and specialization.

Chitinases are ancient enzymes that are widely distributed across all kingdoms of life. Early phylogenetic analyses of fungal GH18 domains proposed a division into three major clades A, B and C, which remains the dominant framework for describing chitinase evolution^25,26^. These classes are distinguished by sequence-level divergence and the presence or absence of a CID. Although these groupings capture some broad trends and appear reasonably robust to taxonomic sampling, subsequent analyses demonstrate they do not consistently correspond to monophyletic clades and decompose into variable, inconsistently defined subclades^27,28^. This has led to conflicting nomenclature, hampering the capacity to systematically link sequence variation to enzymatic activity, ecology or domain architecture across the family. Thus, the evolutionary and functional predictive power of these existing frameworks is limited^27^. Despite extensive individual characterization of diversity across GH18 variants, the question of how domain architecture and insertions/deletions (InDels) affect the evolutionary dynamics of GH18 domain diversification remains unknown.

Enzymes frequently evolve as multi-domain proteins, in which a conserved catalytic core is coupled to auxiliary domains that may tune substrate capture, localization, or regulation^29–32^. Although modularity is widely invoked to explain rapid functional innovation, the rules that govern which domain combinations persist, and how accessory modules co-evolve with catalytic scaffolds, are not fully understood^33,34^. GH18 chitinases are a tractable model to help build our understanding of this process. The family is recognized to encompass architectural diversity and existing work converges on the premise that auxiliary domains are a major axis of chitinase diversification. Moreover, studies have consistently documented changes in activity and substrate preferences upon perturbation of auxiliary domain composition, while domain architecture patterns have been shown to be correlated with function and phylogeny, albeit at restricted, taxonomically constrained scales^27,35^. However, a holistic framework describing underlying mechanisms through which auxiliary domain repertoires may drive GH18 divergence that integrates the observed sequence and structural level diversity of the catalytic TIM-barrel remains elusive. In this work, using family-wide sequence and phylogenetic analyses, we show that auxiliary domain recombination in the GH18 family is non-random and define two complementary axes along which auxiliary domain composition and the catalytic GH18 domain coevolve, driving distinct specialization trajectories. This coordinated modular evolution links sequence architecture and structural changes to selection for niche exploitation and specific physiological roles, providing general principles for how multi-domain enzymes evolve.

## Results

### Domain architecture is a driver of functional divergence in the GH18 family

To map the sequence space of the GH18 family, we curated a comprehensive dataset of 39,582 bacterial and archaeal GH18 sequences (InterPro: IPR001223)^36^. GH18 sequences are defined by a catalytic domain that enables chitin hydrolysis, yet many sequences within this family also contain additional auxiliary domains (e.g., carbohydrate binding). Hereafter, we will refer to GH18 as a family and the various domains as either catalytic (i.e., chitin hydrolysis) or auxiliary (e.g., carbohydrate binding). We first used InterProScan to delineate catalytic and auxiliary domains within each sequence to enable assessment of how domain architecture correlates with GH18 chitinase diversity (Supplementary Note 1, Supplementary Tables 1-5)^37^. While the majority of the GH18 sequences (94.3%) contain a single catalytic domain, we also observe approximately two-thirds of the dataset comprise multi-domain architectures in which a catalytic domain is fused to an auxiliary domain (Figure 1, Supplementary Note 2, Supplementary Figure 1). We identified 110 sequences with more than 8 domains, with a maximum of 18 domains detected for a single sequence (Supplementary Table 6). Previous studies at the kingdom level demonstrate that across prokaryotes, multidomain architectures broadly evolve through random domain accretion, with the number of domains in the fusion protein following an exponential decay law^38,39^. However, here we observe the distribution of total domain counts within a sequence, both catalytic and auxiliary, departed markedly from that expected under random accretion (χ² = 18,891, p < 1×10^−16^), with a strong depletion of single-domain sequences (standardised residual z = −81.1) and pronounced enrichment of multi-domain architectures, particularly three- and four-domain architectures (z = 120.9 and 109.4, respectively; Figure 1b, Supplementary Table 6) ^38^. This is indicative of a non-random organization of domain architectures within the family and suggests three- and four-domain sequences have arisen primarily at the expense of single-domain forms.

**Figure 1.**
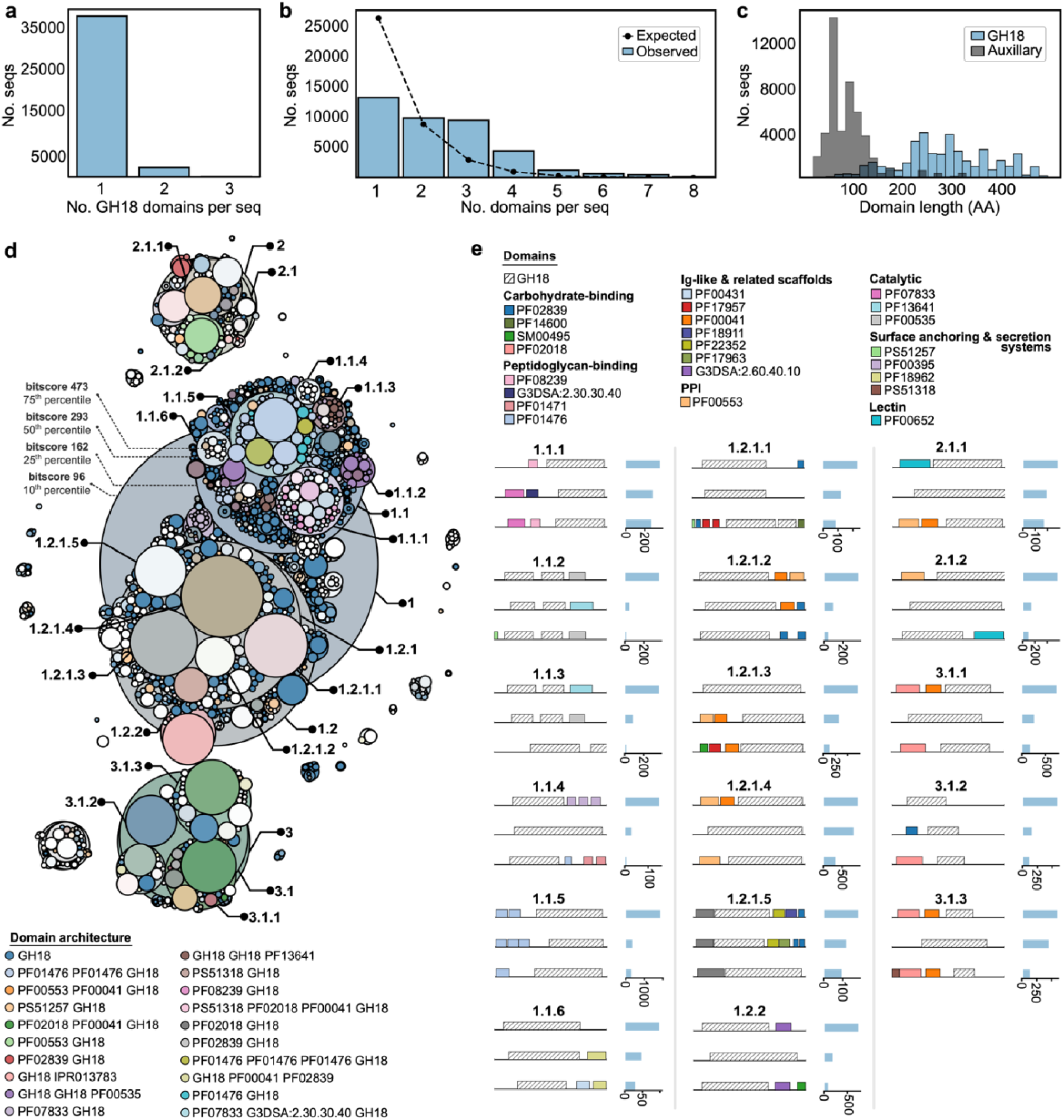
GH18 dataset overview and sequence space visualization. (a) The number of GH18 domains identified within each sequence by InterProScan plotted by frequency. (b) The number of total domains per sequence, as identified by InterProScan is plotted by frequency. The distribution expected under exponential decay (random domain accretion) is plotted as a dotted line. The graph is truncated to sequences with up to 8 domains; although sequences with up to 18 domains were detected, their frequency was so low as to appear negligible at this scale. Complete data displayed in Supplementary Table 6. (c) The distribution of sequence length across auxiliary and GH18 domains. (d) A sequence similarity network of GH18 domains visualized using the ProteinClusterTools workflow^42^. Clusters are coloured by the top 20 most common domain architectures. Key clusters are labelled 1-3, with subclusters indicated. (e) Domain diagrams of the top three most common architectures within key clusters labelled on the SSN. The count of each architecture within each cluster is indicated in the right-hand panel for each domain figure. Sequence length has been standardized for ease of viewing. See Supplementary Table 7 for detailed biological class and annotation descriptions for each domain signature.

The auxiliary domains found in the GH18 family are generally small and span a wide range of functional and structural classes (Figure 1c, Supplementary Table 5 and 7). A total of 365 unique auxiliary domains were detected by InterProScan, which are recombined into 1682 different architectures across the family^37^. The most common auxiliary modules were substrate-binding modules, with the three most abundant being the carbohydrate-binding module family 5/12 (PF02839; 5964 occurrences), a LysM domain associated with general peptidoglycan binding function (PF01476; 5829 occurrences), and a fibronectin type III domain involved in cell adhesion (PF00041; 5563 occurrences; Supplementary Data File 1). The most common multi-domain architecture consists of two tandem LysM domains followed by a catalytic GH18 module (2053; Supplementary Figure 1).

Although the specific mechanisms underlying domain recombination remain unclear, a pronounced size disparity was observed in this dataset with auxiliary domain lengths averaging 76.5 residues (± 48.3) compared to catalytic domains which averaged 270.7 ± 92.5 residues (Figure 1c).This reflects the general observation in the literature that domain mobility appears to negatively correlate with size^40,41^. Broadly, these observations are consistent with a model wherein the catalytic domain is furnished with additional auxiliary domains, as opposed to the catalytic domain functioning as an auxiliary component to a wide variety of unrelated functional cassettes. The pronounced skew toward multidomain architectures, and the corresponding depletion of single-domain forms is indicative of a non-random pattern of domain organization, establishing auxiliary domain diversification as a dominant feature in the evolution and broad functional diversification of the GH18 protein family.

### Conserved domain architecture delineates GH18 domain sequence space

To visualize the sequence space of the GH18 family, we constructed a sequence similarity network (SSN) of only the catalytic domains, using ProteinClusterTools (Figure 1d)^42^. Clustering thresholds were applied at bitscore values corresponding to the 10th, 25th, 50th, and 75th percentiles of the underlying distribution (bitscores of 96, 162, 293, 473, respectively). Three main clusters are observed at the 10th percentile threshold (Clusters 1, 2, 3). Cluster 1 is the largest, comprising 26,945 sequences (68% of the domain dataset analysed) and is further split into two major Clusters: 1.1 and 1.2, with 10,236 and 14,603 sequences, respectively. Clusters 2 (3,363 sequences) and 3 (6,098 sequences) comprise major satellite clusters. Notably, subclusters within 1.1 separate into smaller subclusters at the highest bit score whereas subclusters in Cluster 1.2 remain connected at this stringency. This demonstrates the core catalytic domain sequences within Cluster 1.2 exhibit greater levels of pairwise similarity.

Mapping auxiliary domain architecture to the respective GH18 catalytic domains in the SSN reveals that catalytic domain with equivalent auxiliary domain compositions cluster together (Figure 1e). Clusters are either completely dominated by a single architecture (e.g., 1.1.5, 1.1.2), or alternative arrangements of the same domains and/or a small collection of domains (e.g., 1.1.1, 1.2.1.5, 3.1.3; Figure 1e). This begins to emerge at the 25th percentile threshold and is clear at the 50th percentile threshold, particularly for Cluster 1.1. Qualitatively, this suggests subclusters within Cluster 1.1 comprise more specialized catalytic domains within conserved domain architectures whereas subclusters within Cluster 1.2 comprise catalytic domains of greater sequence similarity, and that these more highly conserved catalytic domains are associated with more diverse combinations of a core set of auxiliary modules. Put simply, there appears to be a dichotomy: members appear to have diversified either through changes to the catalytic domain (with relatively consistent auxiliary domains; Cluster 1.1), or through diversification in the auxiliary domains (with relative conservation of the catalytic domain; Cluster 1.2). Thus, auxiliary domain composition appears to be a key factor in catalytic domain sequence divergence, consistent with co-specialization and cooperative function between neighbouring modules *in vivo*.

### Auxiliary domain promiscuity correlates with GH18 domain sequence conservation

To quantify the disparate evolutionary dynamics governing domain accretion across the GH18 family, we analysed domain combinations across the dataset to distil any patterns associated with domain architecture diversity (Figure 2a). Protein domains vary in their combinatorial potential; some are restricted to specific partners, while others are more versatile and appear in diverse architectures^43–47^. To quantify this, we adapted the Domain Versatility Index (DVI) to calculate a ‘promiscuity’ score (Φ) for each auxiliary domain by comparing neighbour diversity (*T*) to the domain’s total abundance (*n*; see Methods and Supplementary Note 3)^46,48^. This allows for the determination of whether a domain appears in more distinct architectures than expected by chance. To confirm that domains with high promiscuity scores were indeed associated with diverse architectural contexts, we analysed a subset of domains with high (Φ > 0.01), moderate (Φ ∼ 0.01) and architecturally restricted (Φ < 0.008) promiscuity scores (Figure 2b). This confirmed that promiscuous domains, such as the ‘Putative Ig domain’ PF05345, are indeed found in varied architectures with larger numbers of partner domains, moderately scoring domains, such as the ‘Bacterial Ig domain’ PF17936, exist in many ‘pseudo-promiscuous’ variations of similar core architectures, and finally architecturally restricted domains, such as the ‘PE motif family’ domain PF00934, are generally found in highly similar architectures.

**Figure 2.**
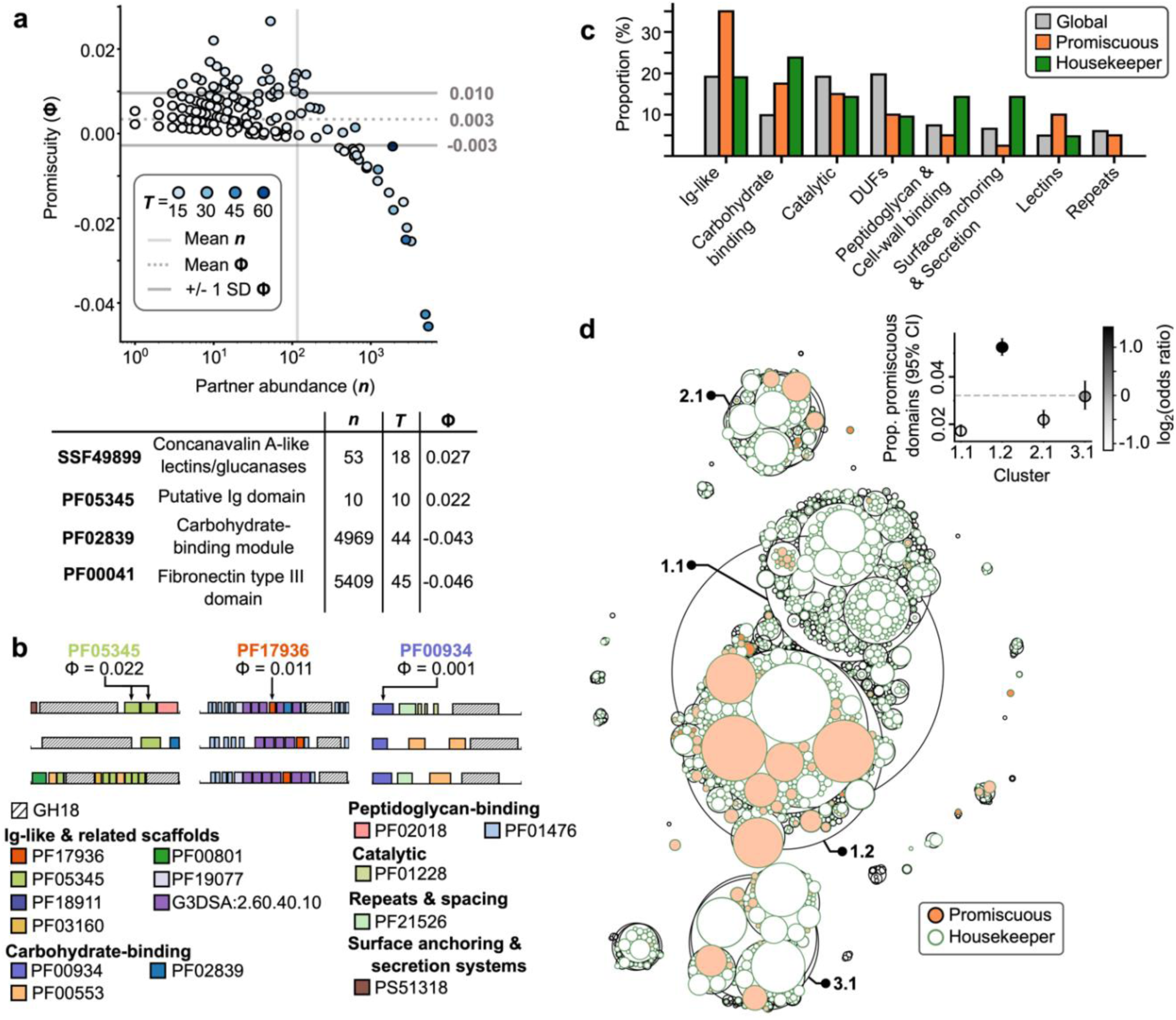
GH18 auxiliary domain promiscuity. (a) Domain abundance (n) is plotted by promiscuity relative to GH18 domains in the complete dataset. Light grey lines at y = 0.010 and y = −0.003 indicate Φ values one standard deviation above and below the mean, respectively. The mean Φ value (y = 0.003) is indicated by a light grey dotted line. The mean of n is indicated by the vertical light grey line at x = 116. Each point is coloured by the number of unique neighbourhoods in which the domain is observed within the dataset (T, legend in figure). (b) Example domain architectures containing common promiscuous domains. (c) The proportion of the complete auxiliary domain dataset, the promiscuous domain dataset (domains with Φ > 0.010) and the housekeeper domain dataset (domains with Φ < −0.003) within each broad biological function category. (d) The SSN presented in Figure 1 with cluster fill coloured depending on the presence of promiscuous domains in the broader GH18 sequence architecture and outlined in green depending on the presence of housekeeper domains. An inset panel displays the proportion of GH18 catalytic domains associated with a promiscuous (Φ > 0.010) auxiliary domain within a given cluster, with vertical error bars showing Wilson’s 95% confidence intervals for the binomial proportion, representing uncertainty in the estimated proportion based on cluster size. Point colour reflects the log₂-transformed odds ratio from Fisher’s exact tests comparing each cluster to the global background frequency, providing an effect-size measure of enrichment or depletion. The horizontal dashed line denotes the dataset-wide background rate of proteins containing at least one promiscuous domain (y = 0.032).

The DVI analysis revealed two outlier classes of auxiliary domains: broadly conserved ‘housekeeper’ domains displaying low promiscuity scores and high abundance values (Φ < −0.003, one standard deviation below the mean), such as carbohydrate binding modules (PF02839) and secretion signals (PF00934), and a subset of highly promiscuous domains with a promiscuity score more than one standard deviation above the mean (Φ > 0.01), consistent with elevated recombination potential, such as concavalin A-like lectins (SSF49899) and Ig domains (PF05345) (Figure 2a,b). Promiscuous domains are enriched in folds associated with adhesion, substrate recognition, and modular recombination, such as Ig-like β-sandwich folds (≈18% vs 8% in the global set), and lectin-type sugar-binding modules (≈5% vs 3%) (Figure 2c, Supplementary Tables 8-10). This overrepresentation suggests that promiscuous domains function as mechanically flexible, combinatorial units that facilitate rapid diversification of binding modalities and substrate engagement/localization. In contrast, low-promiscuity ‘housekeeper’ domains are present in 62% of all sequences and are enriched in carbohydrate-binding domains (23.8% *vs.* 9.9% of the global dataset), surface anchoring/secretion ((14.3% vs 6.6% in the global set) and cell-wall associated domains (14.3% vs 7.4% in the global dataset) (Figure 2c, Supplementary Tables 8-10). These domains also appear in stable, repetitive combinations. Housekeeper domains thus appear to constitute a conserved “structural grammar,” providing the essential scaffolding upon which more complex GH18 architectures can diversify.

Mapping auxiliary domain promiscuity to the GH18 sequence space reveals that promiscuous domains are significantly enriched within Cluster 1.2 and depleted in Cluster 1.1, while housekeeper domains are broadly distributed across all clusters (Figure 2d). These results quantify the qualitative patterns visualized in the SSN (Figure 1). While the catalytic domain sequences within Cluster 1.2 are more conserved, as indicated by large clusters remaining intact at high bit score thresholds in the SSN, they are associated with more diverse domain auxiliary domain arrangements. Comparatively, the more divergent catalytic domains characteristic of Cluster 1.1 have fewer promiscuous domains (Figure 2d). Collectively, these findings further support the observation that GH18 evolution has proceeded along two complementary axes: one in which divergence is driven by external modular recombination, exemplified by sequences within Cluster 1.2, and another wherein specialization of catalytic domains with conserved auxiliary domain architectures drives catalytic domain divergence, as illustrated by sequences within Cluster 1.1.

### Phylogenetic analysis reveals novel sub-families with progressive structural elaboration

To further investigate the relationship between domain accretion and catalytic domain diversification, we inferred a maximum-likelihood phylogeny from 784 GH18 catalytic domains selected to maximize sequence-space coverage and stabilize long branches. Five independent IQTREE3 reconstructions yielded four replicate topologies with consistent placement of five main clades (Figure 3a)^49,50^. A single replicate clustered divergent sequences at the two deepest bifurcations (at the base of Clades I/II and the clade containing III/IV/V), producing a topology that passed the approximately unbiased test but exhibited poor bootstrap support (Supplementary Table 11, Supplementary Figure 3). This replicate was excluded, and the topology shown in a, which exhibited the strongest support at all five defining nodes, was used for downstream analyses. Comparing the phylogenetic tree with the SSN cluster information reveals strong correspondence: Clade I comprises sequences from Cluster 3, Clade II is populated with Cluster 2 sequences, Clade III is comprised of with Cluster 1.2 sequences characterized by high sequence conservation and extensive auxiliary-domain promiscuity, while Clades IV and V correspond to Cluster 1.1 sequences characterized by more divergent GH18 catalytic domain sequences and relatively conserved auxiliary domains.

**Figure 3.**
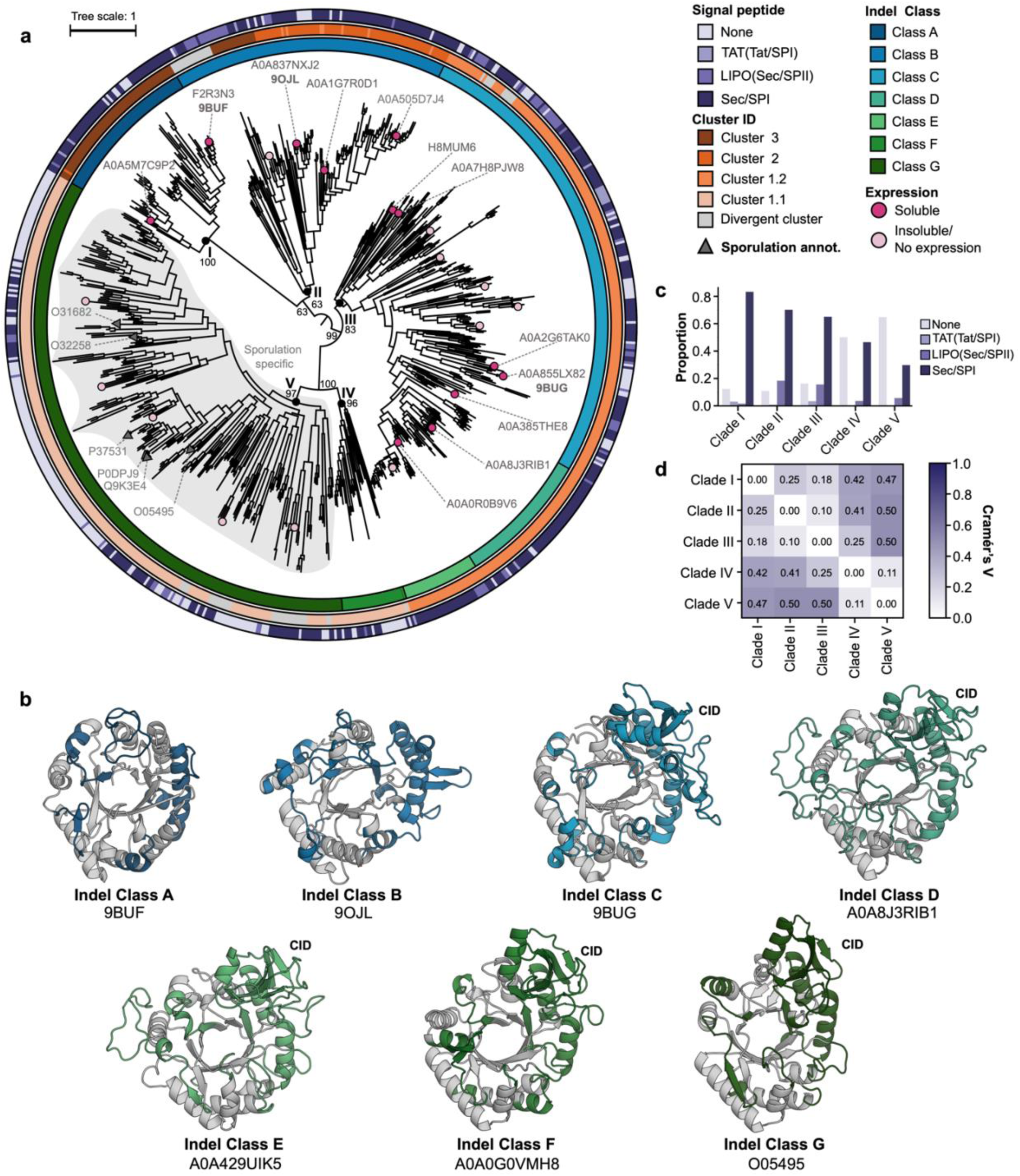
Maximum likelihood phylogeny of representative bacterial GH18 domains. (a) Key clades are labelled at the basal node; bootstrap supports of key clades and other basal bifurcations are labelled. Pink circles indicate leaves selected for expression with light pink indicating no expression/insoluble candidates and dark pink indicating expressed and characterized variants. Grey triangles indicate reviewed sequences annotated in UniProt as sporulation-specific. A grey outline traces the inferred boundary of sporulation-related GH18 domains. From the inside out, colour strip annotations indicate for each sequence: InDel class label, cluster ID and SignalP 6.0 prediction^53^. The tree is midpoint rooted. (b) Crystal structures and AlphaFold models of key CID motif representatives^54^. Regions corresponding to key insertions distinguishing each motif group in the MSA and the chitinase insertion domain (CID) region are coloured as per the key in (a). (c) The proportion of signal peptide categories per clade as predicted by SignalP 6.0^53^. (d) Heatmap detailing the pairwise Cramér’s V score comparing the signal peptide composition of each clade.

Unfortunately, no clear outgroup could be resolved, precluding confident designation of any lineage as a reference and motivating the use of internal rooting criteria (Supplementary Note 4). The phylogeny shown in Figure 3a is therefore presented midpoint-rooted to provide consistent orientation of the maximum likelihood topology. Across all replicates, midpoint-rooting consistently places a deep basal bifurcation near the root of the tree, splitting Clades I and II into a distinct monophyletic group relative to Clades III, IV, and V. Notably, alternative rooting approaches, including Minimal Ancestral Deviation (MAD) rooting and inference under the nonreversible nQ.pfam model, recovered concordant large-scale topology and supported the same overall rooting orientation (Supplementary Figures 3 and 4)^51,52^.

The phylogeny presented here is substantially larger and more complete than those reported previously^25,55^, and challenges the historical A/B/C classifications^6,25,55–57^; accordingly, we propose a revised nomenclature based on this phylogeny and the corresponding InDel architecture. We have identified seven distinct InDel Classes (A-G) distributed non-randomly across Clades I-V (Figure 3), determined based on the presence and sequence signatures of insertions at nine sites within the catalytic domain, including the canonical CID (Table1, Supplementary Figure 5). InDel Classes A and B, found in Clades I and II, respectively, lack the canonical CID and feature minimal insertions, with class A consisting almost entirely of the β-barrel core, and class B exhibiting only short permissive loops. We solved crystal structures of representative proteins from both Clade I (9BUF; 1.75 Å) and Clade II (9OJL; 1.8 Å), revealing that these compact insertions primarily form loops linking the N-terminal of core β-sheets within the catalytic TIM barrel with each other and/or other structural motifs (Figure 3b, Supplementary Table 12). These InDel Classes also possess an extended ɑ-helix that is otherwise truncated in Classes C-G to accommodate the CID motif. The canonical CID appears in Clades III to V, with each InDel Class further distinguished by characteristic loop augmentations (Figure 3a,b). Loop accretion is most pronounced in Clade 3, which exhibits stepwise structural elaboration across InDel classes C, D, and E (Figure 3b). A crystal structure of a Clade III representative confirms this (9BUG; 1.30 Å; Figure 3, Supplementary Table 12). By contrast, InDel Classes F and G, found in Clades IV and V, respectively, are less architecturally ornate but retain the CID. To formalize this evolutionary progression and facilitate future classification, we propose an InDel Class A-G naming scheme (Table 1). This InDel analysis is consistent with the phylogenetic tree and suggests a parsimonious evolutionary scenario in which GH18 variants with fewest insertions (InDel classes A and B, comprising Clades I and II) form a basal monophyletic group (noting the mid-point root), with increasing structural elaboration radiating into the adjacent branch containing Clades III, IV, and V. This trend is visible as a progressive increase in insertional complexity from Clade I to Clade III (InDel classes A through to E), followed by two lineage-specific elaboration patterns in Clades IV and V (InDel classes F and G).

**Table 1.** Detailed description of the sequence and structural sites of the nine insertions used to categorize InDel Classes 1-7. The multiple sequence alignment in question has been included as Supplementary Data File 2 and a detailed description in Supplementary Figure 5.

| Insertion | Position in MSA | Structural site | Extended insertion Class | Moderate insertion Class | Absent Class |
| --- | --- | --- | --- | --- | --- |
| 1 | 8-31 | Loop between β-sheets 1 and 2 | C, D, E, G | A, B, F | |
| 2 | 47-97 | Loop between β-sheet 2 and α-helix 1 | D, E | C | A, B, F, G |
| 3 | 123-130 | Loop between β-sheet 3 and α-helix 2 | C, D, E, F, G | | A, B |
| 4 | 159-174 | loop between α-helix 2 and β-sheet 5 | D | E | A, B, C, F, G |
| 5 | 186-203 | Loop between β-sheet 5 and α-helix 3 | C, D, E | B | A, F, G |
| 6 | 215-225 | Extended α-helix 3 and loop connecting to β-sheet 6 | C, D | A | B, E, F, G |
| 7 | 236-259 | Extended loop region after β-sheet 6 directly preceding CID insertion containing minor helical turn. | A, B, G | E, F | C, D |
| 8 (CID) | 279-457 | Between β-sheet 6 (7 globally) and β-sheet 8 (9 globally) of core TIM barrel. | C, D, E, F, G | | A, B |
| 9 | 476-494 | C-terminal loop directly preceding the final α-helix | A | | B, C, D, E, F, G |

### Secretion mode analysis identifies distinct functional specialization across clades

Signal peptides (SPs) are key determinants of GH18 physiological function and, given the predominance of extracellular chitinase activity, are often associated with these sequences^23^. To determine SP presence and type across the GH18 family, all sequences were analysed with SignalP6.0^53^. Among predicted SPs, most were canonical Sec/SPI sequences, consistent with secretion into the periplasm or extracellular milieu (Figure 3b). A subset possessed lipoprotein SPs (LIPO (Sec/SPII)) that direct membrane anchoring, and a smaller subset exhibited TAT-type SPs, characteristic of proteins that fold or incorporate cofactors prior to export (Figure 3b)^53,59^. Projecting secretion profiles onto the phylogeny indicated pronounced differences in secretion preferences between clades. Global χ² tests confirmed that secretion state varies non-randomly across the tree, with Clade V driving the strongest deviation owing to its pronounced enrichment for non-secreted members (p ≈ 2.7 × 10^−40^; Figure 3a). To complement this statistical analysis with an assessment of effect size, we performed pairwise Cramér’s V analyses. These reinforced this signal, revealing very strong effect sizes between Clade V and Clades I, II, and III (V ≈ 0.47–0.50; Figure 3c), consistent with Clade V forming a divergent functional subfamily characterized by intracellular localization, in line with sporulation-linked annotations for several members (Figure 3a). Clade IV showed moderate divergence from Clades I and II (V ≈ 0.41–0.43) and weaker separation from Clade III (V ≈ 0.25), despite a non-significant global χ² statistic (p ≈ 0.18), suggesting an intermediate or structurally distinct niche. Its limited divergence from Clade V (V ≈ 0.11) likely reflects shared tendencies toward reduced secretion, evident in the phylogenetic annotation mappings, but differing balances of Sec/SPI versus non-signal-peptide profiles (Figure 3). Among the major non-sporulation clades, Clades I to III exhibited only low-to-moderate pairwise differences (V ≈ 0.10–0.25), forming a continuum from predominantly cytosolic/periplasmic (Clade I) to increasingly secreted (Clades II and III), consistent with functional diversification within a shared cell-envelope context rather than the deep ecological separation observed in Clade V.

### The substrate preference of isolated GH18 domains is relatively broad

To evaluate the catalytic specificity across the GH18 family, we heterologously expressed 25 GH18 catalytic domains in *Escherichia coli* BL21(DE3) (SI Table 13; SI Figure 6). Notably, no Clade V enzymes could be produced in soluble form (Supplementary Figure 6, Supplementary Table 13). This may reflect a requirement for specialized localization, or co-expression with certain auxiliary domains, for correct folding. In contrast, 11 catalytic domains from Clades I to III were purified and exhibited broadly consistent catalytic profiles (Table 2). Enzymes from Clades I and II showed comparable activity profiles, with mixed endo- and exo-chitinase activity and a slight bias toward the exo-mode. Clade III members typically displayed higher turnover rates and a more heterogeneous substrate distribution: while most retained the canonical exo-mode preference, a greater fraction exhibited near-equivalent activities, and individual enzymes deviated from this pattern (Table 2). Notably, A0A0R0B9V6 from Clade III showed disproportionately high endo-chitinase activity, whereas A0A855LX82, also from Clade III, was the only enzyme to display substantial chitobiosidase activity, producing GlcNAc dimers rather than monomers (Table 2). These results are broadly consistent with Clade III GH18 members exhibiting a greater diversity of catalytic specificity.

**Table 2.** Activity profiles of characterized GH18 sequences. Exo-chitinase (4MU-N-acetyl-β-D-glucosaminide), endo-chitinase (4MU-β-D-N,N′,N′′-triacetylchitotriose) and chitobiosidase (4MU-N,N′-diacetyl-β-D-chitobioside) activity are reported for each sequence, the substrates used to test these activities are reported in italics for each respective column. The ratio of average exo- to endo-chitinase activity is also reported.

| Uniprot ID | Clade | InDel Class | Organism | Native domain architecture | Expressed architecture | Activity (μmol/min) |  |  | Exo:endo-activity ratio |
| --- | --- | --- | --- | --- | --- | --- | --- | --- | --- |
|  |  |  |  |  |  | Exochitinase | Chitobiosidase | Endochitinase |  |
| A0A5M7C9P2 | I | A | *Saccharopolyspora hirsuta* | PS51318<br>GH18 | GH18 | 0.995±0.155 | 0.006±0.001 | 1.02±0.02 | 0.98 |
| F2R3N3 | I | A | *Streptomyces venezuelae* | PF00553<br>GH18 | GH18 | 8.62±0.93 | 0.368±0.159 | 4.53±0.15 | 1.90 |
|  |  |  |  |  | PF00553<br>GH18 | 3.17±0.14 | 0.00583±0.0021<br>3 | 1.72±0.12 | 1.84 |
| A0A1G7R0D1 | II | B | *Chitinophaga filiformis* | GH18<br>PF17957<br>GH18<br>PF17957<br>PF02839<br>PF17957<br>SM00495 | GH18<br>*Data from second GH18 domain* | 4.26±0.14 | 0.004±0.001 | 2.71±0.03 | 1.57 |
| A0A505D7J4 | II | B | *Streptomyces sporangiiformans* | PF02018<br>PF00041<br>GH18 | GH18 | 5.34±0.13 | 0.00563±0.002 | 2.24±0.09 | 2.38 |
| A0A837NXJ2 | II | B | *Vibrio splendidus* | GH18<br>PF02839<br>PF21868<br>SM00495 | GH18 | 3.54±0.07 | 0.454±0.784 | 1.91±0.05 | 1.85 |
| A0A0R0B9V6 | III | D | *Stenotrophomonas maltophilia* | SM00495<br>PF17957<br>PF00041<br>GH18 | GH18 | 12.1±0.2 | 0.0698±0.008 | 14.6±1.3 | 0.83 |
|  |  |  |  |  | SM00495<br>PF17957<br>PF00041<br>GH18 | 7.10±0.11 | 0.0508±0.007 | 17.8±0.7 | 0.40 |
| A0A2G6TAK0 | III | C | *Flavobacterium sp. 9* | GH18 | GH18 | 4.77±0.26 | 0.251±0.015 | 1.12±0.06 | 4.26 |
| A0A855LX82 | III | C | *Enterobacter cloacae complex sp. ECNIH14* | GH18 | GH18 | 9.23±0.18 | 5.01±0.09 | 7.05±0.32 | 1.31 |
| A0A8J3RIB1 | III | D | *Sphaerimonospora thailandensis* | PF00553<br>PF00553<br>PF00041<br>GH18 | GH18 | 1.98±0.15 | 0.0156±0.0025 | 2.10±0.15 | 0.94 |
| H8MUM6 | III | C | *Corallococcus coralloides* | G3DSA:2.60.120.430<br>G3DSA:2.60.40.10<br>SSF48726<br>GH18 | GH18 | 4.87±0.29 | 0.0087±0.0068 | 2.45±0.25 | 1.99 |
| A0A7H8PJW8 | III | C | *Aquimarina sp. TRL1* | SM00495<br>GH18<br>PF17957<br>PF17957<br>SM00495<br>PF02839<br>PF18962 | GH18 | 18.2±0.5 | 0.0119±0.0102 | 10.1 ±0.3 | 1.80 |

We also examined the effect of auxiliary domains, confirming that they exert measurable, substrate-dependent effects on catalysis (Table 2). Removal of the N-terminal cellulose binding domain (PF00553) from F2R3N3 (Clade I) enhanced both endo- and exo-chitinase activities, consistent with substrate sequestration by CBM2 when assayed with short fluorogenic substrates. Conversely, inclusion of the three N-terminal auxiliary domains (a chitin binding domain (SM00495), a bacterial Ig domain (PF17957) and a Fibronectin type III domain (PF00041)) in A0A0R0B9V6 increased endo-activity relative to the isolated catalytic core. This confirms that the presence of auxiliary domains can modulate substrate engagement or access to the catalytic cleft. Together, these data are consistent with previous studies, confirming that auxiliary domains are active determinants of GH18 catalytic specificity, tuning substrate presentation, shaping endo/exo balance, and in some clades potentially functioning as architectural prerequisites for proper folding or secretion^60–62^. This highlights a modular evolutionary framework in which domain accretion contributes directly to enzymatic diversification within the GH18 family.

### Taxonomic enrichment and functional annotation contribute to an evolutionary model of chitinase divergence and specialization

To connect the quantitative bioinformatic analyses to ecology and physiology, the SSN (clusters) and phylogenetic (clades) GH18 classifications were overlaid with secretion profiles, taxonomic distributions, and functional annotations from literature. These data resolve a hierarchical model of GH18 evolution in which catalytic domain divergence and auxiliary domain accretion co-evolve along distinct axes to facilitate ecological exploitation and physiological specialization with characteristic combinations of InDel class, auxiliary-domain architecture and secretion mode (Figure 4).

**Figure 4.**
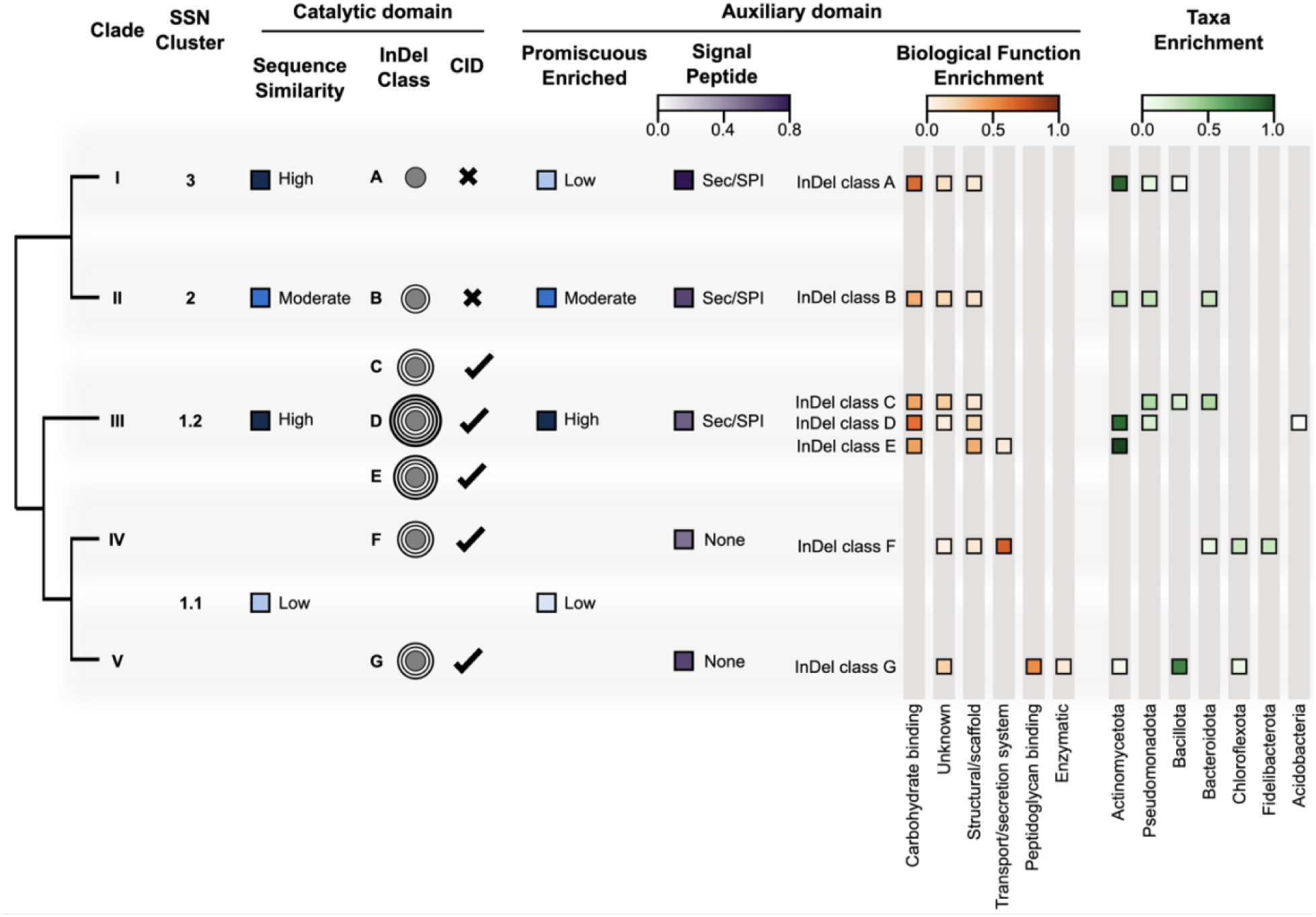
The modular evolution of GH18 chitinases A schematic overview of the five major GH18 clades that resolves the relationship between catalytic domain evolution, auxiliary domain accretion, and ecological function. The figure aligns phylogenetic Clades (I to V) with their corresponding SSN Clusters (1.1, 1.2, 2, 3). Key catalytic domain annotations for each clade are included, incorporating sequence similarity information from the SSN, InDel class A to G with increasing loop elaborations indicated by a circular graphic where increasing rings represent increasing presence/complexity of loop regions, and the presence of the Chitinase Insertion Domain (CID) is also indicated. Auxiliary domain information summarizes the enrichment of promiscuous domains and the dominant signal peptide types. The top three most common functional categories of auxiliary domains for each clade (e.g., carbohydrate binding vs. cell wall anchoring) are displayed as a heatmap with the colour indicating the proportion of sequences in the clade that contain an auxiliary domain associated with that category. Finally, the taxonomic enrichment of each InDel class is also displayed via a heatmap indicating the abundance of specific phyla (e.g., Actinomycetota, Bacillota) within each class. All heatmap enrichment calculations were performed from results on the global GH18 protein family dataset.

We propose Clades I (Cluster 3) and II (Cluster 2), comprising InDel Classes A and B, represent the most simple, ancestral-like states of the GH18 family, reminiscent of what was recently reported for the GH19 family^63^. These enzymes lack the canonical CID and possess minimal insertions, with catalytic domains typically associated with signal peptides and enriched in carbohydrate-binding auxiliary domains, consistent with broad-spectrum extracellular carbohydrate turnover (Figure 3, Figure 4, Supplementary Figure 8, Supplementary Table 16). Despite their similar architectures, the two clades show contrasting taxonomic and modular patterns. Clade I is largely restricted to Actinomycetota, exhibits high catalytic sequence conservation, and is strongly enriched in non-promiscuous, carbohydrate-binding auxiliary domains, consistent with retention in chitin-rich soil environments (Figure 4, Supplementary Table 14-16, Supplementary Figures 7-9). In contrast, Clade II spans Actinomycetota, Pseudomonadota and Bacteroidota and shows a higher incidence of promiscuous auxiliary modules, suggesting a more diffuse lineage with greater modular flexibility, more reminiscent of a basal lineage, and a higher degree of auxiliary domain promiscuity (Figure 4, Supplementary Table 14-16, Supplementary Figures 7-9).

A major evolutionary transition occurs with the emergence of Clade III (Cluster 1.2), IV and V (Cluster 1.1). Clade III (Cluster 1.2) is comprised of archetypal secreted chitinases in which a conserved catalytic core sequence is progressively elaborated by loop insertions, corresponding to InDel Classes C, D and E (Figure 3, Figure 4). Additionally, sequences in this clade are significantly enriched in promiscuous auxiliary domains relative to other clades, most notably carbohydrate binding modules and structural scaffold-like domains (Figure 2, Figure 4, Supplementary Figures 8 and 9, Supplementary Table 16). These features are consistent with fine-grained tuning of substrate presentation and localization atop a conserved chitinase core. This aligns with the taxonomic distribution of the clade, which is skewed towards Actinomycetota, Bacillota and Pseudomonodota; phyla found in glycan-rich niches from soil, marine particle-associated communities, gut microbiomes and plant-litter ecosystems^64,64,65 66^ (Figure 4, Supplementary Figure 7, Supplementary Tables 14 and 15). As sequences radiate into InDel classes D and E, subclades become dominated by Actinomycetota, particularly with *Streptomyces* sequences. Sequences included in the phylogeny were selected non-arbitrarily from the GH18 protein family using the ProteinClusterTools workflow to capture sequence-level diversity independent of taxonomy^42^. That these taxonomically biased clades are robust to this sampling method indicates that substantial GH18 radiation within these lineages relative to the global family, consistent with intense niche specialization and exploitation. Altogether, specialization within Clade III appears to driven by the coordinated elaboration of loop insertions atop a well conserved catalytic TIM-barrel, together with highly modular domain architecture recombination.

In contrast, the evolution of Clades IV and V (Cluster 1.1; InDel classes F and G) reflects a stepwise exaptation toward lineage-specific physiological roles. These sequences exhibit moderate insertional elaboration and relatively conserved domain architectures but notable sequence-level divergence of the catalytic TIM-barrel and depletion of promiscuous auxiliary modules (Figure 1, Figure 2, Figure 3). Signal peptides are also strongly depleted, indicating chitinases in these clades are not secreted (Figure 3, Figure 4). Clade IV is enriched in transport and membrane-associated auxiliary domains and occurs predominantly in biofilm-associated phyla such as Chloroflexota and Bacteroidota, suggesting periplasmic or cell envelope functions, potentially in local glycan processing or biofilm maintenance (Figure 4, Supplementary Figures 7-9, Supplementary Tables 14-16)^67–70^. Clade V is strongly enriched in spore-forming Firmicutes (Bacillaceae, Paenibacillaceae) and in peptidoglycan-binding auxiliary domains, and includes multiple proteins annotated as sporulation-specific glycosylhydrolases (Figure 3, Figure 4, Supplementary Figures 7-9, Supplementary Tables 14-16)^64,71–76^. These features support a model in which GH18 domains have been functionally exapted as sensor–effector components in cortex remodelling and germination^77–80^.

## DISCUSSION

The systematic exploration of bacterial GH18 sequence space presented here unifies previously disparate observations on chitinase diversity into a single evolutionary framework. By integrating auxiliary-domain architecture, internal insertion/deletion patterns and phylogeny, we resolve the family into five main clades that are better described by a graded series of loop architectures (InDel classes A–G) than by the historical A/B/C categories^8,56,81^. These classes capture a stepwise accretion and reshaping of internal loops, including the canonical CID, and can be assigned using a simple HMM-based scheme that operates directly on sequence^82^. The resulting map provides a structurally interpretable model for GH18 evolution: closely related InDel classes occupy neighbouring regions of sequence and structure space, yet differ systematically in their secretion mode, auxiliary-domain usage and ecological contexts. In this view, GH18 diversification is governed by the co-evolution of a conserved TIM-barrel scaffold, its internal loop architecture and a restricted set of auxiliary modules, rather than by unconstrained domain shuffling around an invariant catalytic core.

The domain-architecture analyses reveal a non-random modular grammar of assembly that underpins how auxiliary modules are combined with catalytic domains. Instead of following the near-exponential decay of domain counts reported at the proteome scale, GH18 architectures are skewed towards multi-domain forms, with three-domain configurations strongly over-represented (Figure 1b)^38,39^. Quantifying these dynamics through a DVI-based analysis resolved auxiliary domains into distinct classes. While most appear as rare specialists that recombine as expected given their abundance, subpopulations of (i) ‘promiscuous’ domains associated with exceptionally diverse architectures, and (ii) highly abundant but architecturally conservative ‘housekeeper’ domains are evident (Figure 2). These housekeeper domains thus form static architectural backbones that are heavily exploited but rarely recombined into novel configurations, while promiscuous domains appear to enable architectural recombination. These findings recapitulate, within a single ancient enzyme family, the hub-and-spoke organization of domain space described at the proteome level, and shows directly how a small set of static and mobile modules jointly constrain and enable exploration of GH18 modular space^31,33,83^. Additionally, these labels are associated with strong context-dependence, with domains such as the carbohydrate binding domain CBM14, which have been shown to be globally promiscuous, behaving as housekeeper modules within the GH18 neighbourhood^84^. This indicates that domain mobility may be an emergent property of the neighbouring domains, rather than an intrinsic feature of particular folds^46,48^.

This work clarifies two complementary diversification strategies in GH18 evolution that correspond to distinct ecological roles. Clade III (Cluster 1.2) exemplifies a ‘modular scavenger’ trajectory distributed across Actinomycetota, Bacillota and Pseudomonadota, phyla that dominate in glycan-rich environments such as soil, marine particles and gut communities. Here, we propose the catalytic TIM-barrel remains highly conserved while internal loops are elaborated, and auxiliary domains readily recombined to jointly tune substrate presentation, processivity and localisation against a conserved chemical core. How activity is modulated via remote loop accretion over evolution is not fully understood in GH18 chitinases, although the capacity for such features to alter activity has been documented in GH19 enzymes^63^. Equally, auxiliary context and cellular localisation have been shown here to enable fine-grained functional specialisation *in vivo* without altering the core (Table 2). In this ecological context, these modes of expansion may be favoured to promote capture and processing of heterogeneous, recalcitrant chitin substrates while preserving catalytic robustness across variable external conditions^14,85,86^. In the second trajectory, represented by Clades IV and V, catalytic sequences diverge more markedly, loop elaboration is modest, and auxiliary architectures are relatively fixed and enriched for peptidoglycan-binding and membrane-associated functions, with a pronounced loss of secretion signals. Here, GH18 domains appear to fulfil extracellular nutrient scavenging towards specialized roles in the cell envelope and developmental transitional such as sporulation, where stringent control over regulation and localization are likely to be more important than broad substrate range^8,24,87,88^. Early, relatively simple clades (I and II) lie at the base of these trajectories, retaining compact cores with limited insertions and broadly distributed, carbohydrate-binding-dominated auxiliary sets, consistent with ancestral secreted chitinases that pre-date the emergence of highly elaborated scavenger and exapted developmental lineages.

More broadly, these findings provide a mechanistic model for how multi-domain enzymes diversify. Rather than treating catalytic domains and accessory modules as independent evolutionary units, the GH18 family shows that catalytic domain sequence, internal loop architecture, auxiliary-domain grammar and cellular localization evolve as a coupled system^31–33,89^. The same catalytic scaffold can support subtly distinct ecological strategies under different modular contexts, while divergent cores embedded in conserved auxiliary backbones adopt more distinct, systematic shifts in function. These principles are likely to extend to other modular systems such as kinases, non-ribosomal peptide synthetases and other CAZymes, where small domains act as recombinable units^46,83^. They also have implications for protein engineering and for data-driven models of protein function. Current language-and diffusion-based methods predominantly optimize isolated domains, whereas natural selection has tuned integrated assemblies in which loop evolution and modular context are inseparable^90^. Incorporating modular design rules, such as the separation of housekeeper versus promiscuous modules, the limited alphabet of viable architectures, and the use of loop accretion as an orthogonal evolutionary mechanism, should improve our ability to design synthetic multi-domain enzymes that retain evolvability and context-appropriate deployment.

## METHODS

### Dataset collection and domain annotation

All protein sequences associated with the bacterial and archaeal chitinase GH18 family (IPR001223; 39,582 sequences) were retrieved from UniProt on October 7, 2025, and their domain architectures annotated using InterProScan (version 6.0.0). Each functional signature was parsed to eliminate annotation redundancy, eliminating equivalent and overlapping functional signatures. Annotations were initially categorized as GH18-related or auxiliary. GH18-related annotations were iteratively clustered on a per-sequence basis whenever coordinate intervals overlapped by more than 20%, and clusters containing at least one GH18 signature were collapsed into a single representative record following the priority rule below, yielding the preferred GH18 domain coordinates for each protein. The consolidation algorithm assigned a priority rank to the three canonical identifiers (PF00704, PS51910, SSF51445, ranked 1, 2, and 3, respectively); any domain in the GH18 signature database that was not associated with one of these top three signatures was assigned the most common signature based on pre-computed global frequencies. All sequences were found to contain at least one domain associated with a GH18 signature. Similarly, auxiliary annotations were subject to a coordinate-based clustering to reduce redundancy, where two annotations were considered the same domain if their start/end coordinates fell within ± 5 amino acids or if the overlap between their intervals was 20% of the shorter span. Connected components in the resulting overlap graph were collapsed to a single representative chosen by a hierarchical rule set: choosing the most common PFAM entries as a priority and then the most globally frequent signature among remaining candidates. Additional notes in Supplementary Note 1 and data in Supplementary Tables 1-5, Supplementary Figure 1.

### Sequence similarity network creation and annotation

The GH18 domains of all sequences in the dataset were extracted and used to generate a homology-based sequence similarity network (SSN). All-by-all pairwise comparisons were performed using MMSeqs easy-search on default settings. Bit scores were used as the metric for homology rather than E-values to maintain resolution at high similarity thresholds and minimize size effects. Network construction, visualization and annotation was performed using the ProteinClusterTools package^42^.

### Domain promiscuity analyses

Domain promiscuity within the GH18 protein family was calculated by adapting the Domain Versatility Index method outlined by Weiner et al. (2008). For each non-GH18 domain, two properties were quantified: (i) how often it co-occurred with GH18 in a protein, and (ii) how many distinct domain neighbours it exhibited within GH18-containing architectures. To count co-occurrence, the set of distinct domains present in each sequence’s architecture was first constructed using the InterProScan annotation workflow outlined above. For each domain *d* in this set, we incremented a co-occurrence count *n*_(GH18, *d*)_ by one. This ensures that *n*_(GH18, *d*)_ reflects the number of GH18-containing proteins in which *d* appears at least once, independent of its multiplicity within a single architecture.

To quantify neighbour diversity, we treated the ordered domain list of each GH18-containing protein as a linear architecture and recorded, for every non-GH18 domain *d* in that list, its immediate left and right neighbours (where present). For each appearance of *d*, the domains at positions *i − 1* and *i + 1* were considered candidate neighbours; if a neighbour existed and was not identical to *d*, it was added to the neighbour set of *d*. Across all proteins, this yielded, for each domain *d*, a set of distinct neighbours, whose cardinality defined the total neighbour diversity *T*_(GH18, d)_ (i.e. the number of unique domains observed adjacent to *d* within GH18-containing architectures). To convert these counts into relative measures, the neighbour diversity of each domain *d* was normalized by the total neighbour diversity summed across all domains in the dataset. Specifically, we defined:

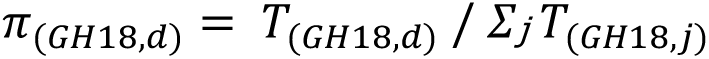

where indexes all non-GH18 domains considered in the analysis (i.e., the same domain set to which *d* belongs). In this formulation, *d* refers to the focal domain for which the normalized diversity is being calculated, whereas *j* ranges over every domain in the dataset to provide the denominator representing the total neighbour diversity. In parallel, the co-occurrence frequency of each domain was normalized by the total number of GH18-containing proteins *f*_(GH18, *d*)_, reflecting how common *d* is among GH18 architectures.

To obtain a GH18-specific promiscuity score (φ), the neighbour diversity of each domain was scaled to its abundance as follows:

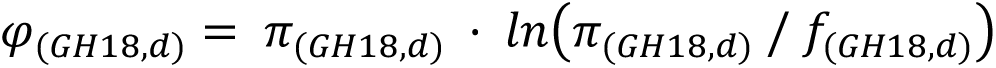

This measure assigns higher scores to domains that, relative to how frequently they appear in GH18-containing proteins, are associated with a disproportionately diverse set of neighbouring domains, and therefore are more likely to participate in a wide range of GH18 architectures.

To evaluate the distribution of promiscuous domains across the SSN, enrichment analyses were performed using Fisher’s exact test (one-sided) on 2×2 contingency tables, comparing category frequencies between specific domain sets and the global annotated domain set. Resulting *p-*values were corrected for multiple testing using the Benjamini–Hochberg False Discovery Rate (FDR) method. All analyses were performed in Python using SciPy and statsmodels. Additional notes and data attached as Supplementary Note 3, Supplementary Data Files and Supplementary Tables 8-10.

### Signal peptide annotation

Signal peptides were predicted for all full-length chitinase sequences using SignalP 6.0.^53^ The software was downloaded and installed locally and predictions were performed on the complete GH18 dataset. The resulting signal peptide and cleavage site predictions were subsequently integrated into downstream analyses for comprehensive sequence annotation.

### Phylogenetic inference

GH18 domain sequences were non-arbitrarily selected from the family through the HMM-based representative selection method implemented through ProteinClusterTools^42^. Representative sequences were selected from every cluster within the SSN across bitscore thresholds from 400 to 800 at increments of 25, resulting in 623 sequences. This dataset was supplemented with all annotated sequences (denoted ‘Reviewed’ on UniProt) to enable downstream tree annotation. Sequences were aligned using MAFFT with the -dash and -localpair settings^91^. Alignments were manually curated and initial trees visualized using FastTree^92^. We attempted to supplement sparse clades and long branches with targeted BLAST searches of the NCBI nr clustered database. Maximum likelihood phylogenies were inferred from a final dataset of 771 sequences using IQTREE3, employing ModelFinder Plus (MFP) to automatically select the best-fitting substitution model (in this case all replicates converged on Q.PFAM+F+R10), with branch support assessed using 1,000 ultrafast bootstrap replicates and the subsequent tree search optimized by best-of-five searches, including BNNI optimization^49–51,93–95^.

### Protein Expression and Purification

Genes encoding selected chitinases were codon-optimized for expression in *Escherichia coli*, synthesized and cloned into pET-29b(+) vectors (adding C-terminal hexahistidine purification tags) (Twist Biosciences). Each plasmid was transformed into chemically-competent *E. coli* BL21(DE3) cells (NEB) using the heat shock and plated onto LB agar supplemented with 50 μg mL^−1^ kanamycin before being incubated overnight at 37 °C. Individual colonies were picked to inoculate 1 mL LB cultures with 50 μg mL^−1^ kanamycin in a 96-well plate. The culture was incubated at 37 °C at 1,050 rpm on a plate shaker for 12 hours. Glycerol stocks were prepared by the addition of 1 ml of 50% glycerol to each culture and stored at −80 °C.

All sequences were initially expressed at small scale in a 2 mL deep 96-well plate. 1.5 mL cultures of simple autoinduction media (5 g L^−1^ Yeast Extract, 20 g L^−1^ Tryptone, 5 g L^−1^ NaCl, 6 g L ^−1^ Na_2_HPO4.7H_2_O, 3 g L^−1^ KH_2_PO_4_, 0.19 g L^−1^ MgCl_2_, 6 g L^−1^ glycerol, 0.5 g L^−1^ glucose, 2 g L^−1^ D-lactose) supplemented with 100 μg mL^−1^ kanamycin were inoculated from glycerol stocks. The 96-well plate was incubated for 24 hours at 37 °C on a plate shaker shaking at 1,050 rpm. Cultures were harvested by centrifugation at 2250 × *g* for 45 min at room temperature. Cell pellets were stored at −20 °C until purification. For large-scale expression in *E. coli* glycerol stocks were used to inoculate a seed culture of 10 mL LB media with 50 μg mL^−1^ kanamycin. The seed culture was incubated at 37 °C for 12 hours shaking at 250 rpm. Seed cultures were then diluted 1:100 to inoculate 200 mL or 1 L culture in simple autoinduction media supplemented with 100 μg mL^−1^ kanamycin in 500 mL or 2 L Thomson Ultra Yield Flasks. The cultures were incubated for 24 hours at 37°C shaking at 250 rpm. Cells were harvested by centrifugation at 4730 × *g* for 15 minutes at room temperature. Cells were frozen at −20 °C until purification.

For small scale cultures, proteins were purified using 96-well plate immobilized metal affinity chromatography (IMAC). Cell pellets were resuspended and lysed in 50 μL BugBuster reagent (Sigma Aldrich) dissolved in equilibration buffer and shaken at 100 rpm for 30 minutes. Insoluble cell debris was separated by centrifugation at 3,000 × *g* for 45 minutes. The supernatant was transferred to a new 300 μL 96-well plate. Insoluble pellets were stored at −20 °C until inclusion body purification with BugBuster reagent (Sigma Aldrich) and analysis by SDS-PAGE. 96-well Hispur Ni-NTA (Thermo Fisher Scientific) spin plates were equilibrated using 200 μL equilibration buffer per well (20mM Tris, 500mM NaCl, pH 7.4). Next, 200 μL equilibration buffer was added and plates were spun at 1,000 × *g* for 1 minute. This was repeated 3 times. The supernatant was transferred to the Hispur spin plate and washed with an equal volume of binding buffer (20mM Tris, 500mM NaCl, 20mM imidazole, pH 7.4) 3 times with spins at 1,000 × *g* for 1 minute between washes. Proteins were eluted by the addition of 200 μL of elution buffer (20mM Tris, 500mM NaCl, 500mM imidazole, pH 7.4) per well to the spin plate and spun at 1,000 × *g* for 1 minute. Eluted proteins were stored at 4 °C until further analysis. Small scale cultures were used for all activity assays.

For large scale cultures proteins were purified using immobilized metal affinity chromatography. Cell pellets were resuspended in 40 mL equilibration buffer and lysed by sonication (2 x 5 minutes at 50% power). Insoluble cell debris was removed by centrifugation at 17400 × *g* for 60 minutes at 4 °C. Supernatants were filtered through 0.45 μm filters before loading onto a 5 mL HisTrap HP (Cytiva) column. Columns were pre-equilibrated with 5 column volumes (CVs) of binding buffer. The column was washed with 15 CVs binding buffer, followed by elution using an increasing gradient of elution buffer from 0-100% over 20 CVs. SDS-PAGE was used to confirm which fractions contained the target protein. Selected fractions were then buffer exchanged into a suitable buffer by dialysis using SnakeSkin Dialysis Tubing, 10K MWCO, 35 mm dry I.D. (ThermoFisher Scientific) overnight at 4 °C.

To purify proteins for crystallization, selected variants were concentrated using an Amicon Ultra-15 10K MWCO Centrifugal Filter Unit (Merck Millipore) before loading onto a size exclusion column (HiLoad Superdex 26/600 200 prep grade, GE Healthcare) pre-equilibrated with 2 CVs size exclusion buffer (20mM HEPES, 150mM NaCl, pH 7.4). Proteins were eluted with an isocratic elution of 2 CVs of SEC buffer. Selected fractions were analysed for purity by SDS-PAGE. Pure samples were pooled and stored at 4 °C until crystallization.

### Protein Crystallization

Proteins were concentrated using a 10 kDa MWCO centrifugal concentrator prior to sparse matrix screening of crystallization conditions using 0.5 µl protein and 0.5 µl well solution in a sitting drop format at 18°C. For A0A855LX82, protein was concentrated to 140 mg/ml and formed diffraction-quality crystals in well A12 of ShotGun1 crystal screen (Mitegen); 20% PEG 8000, 0.1M Sodium cacodylate, 0.2M Magnesium acetate. Crystals were harvested and cryo-cooled without further cryoprotection. For F2R3N3, protein was concentrated to 100 mg/ml and formed diffraction-quality crystals in well D9 of ShotGun1 crystal screen (Mitegen); 30% PEG 2000 MME, 0.2M Ammonium sulphate, 0.1M Sodium acetate. Crystals were harvested and cryo-cooled without further cryoprotection. For A0A837NXJ2, protein was concentrated to 50mg/ml, and sparse matrix screening yielded crystals that required optimization. Following optimization of crystallization conditions, diffraction quality crystals were obtained in a hanging drop format against a well solution of 1.7 M Ammonium sulfate, 0.1 M Bis-Tris pH 7.0, and 0.1M NaCl. Crystals were cryo-protected in well solution with 20% ethylene glycol prior to flash cooling in liquid nitrogen.

### Structure Determination

360° of diffraction data were collected for each protein on the MX2 beamline at the Australian Synchrotron^96^. Data were collected with 80% attenuation of the beam for F2R3N3, with a detector distance of 180 mm. For A0A855LX82, data were collected with 95% attenuation of the beam and a detector distance of 150 mm. Data for A0A837NXJ2 crystals were collected with 80% attenuation and a detector distance of 200 mm. Data were processed using XDS^97^ and truncated using AIMLESS The structures were solved by molecular replacement (Phaser MR^98^, CCP4 suite) using the AF2-predicted models of the corresponding protein GH18 domain as the search models. Refinement was carried out in phenix.refine^99^. Multiple rounds of refinement and model building in Coot v1.1.17^100^ were carried out until R factors converged, and the structure was modelled as best as possible to the electron density. MolProbity was used for structure validation^101^. Structures have been deposited to the Protein Data Bank under accession codes 9BUG, 9BUF, and 9OJL.

### Chitinase activity assays

A fluorescent chitinase assay kit was obtained from Sigma Aldrich (CS1030). For assays on the combined three substrates 10 μL of sample was added to 90 μL of substrate diluted in the provided assay buffer at the recommended concentration in a 300 μL black 96 well plate. For individual assays, substrates were diluted with assay buffer to 320 μM. Samples were incubated for 1 hour at 37 °C, after 1 hour the reaction was halted by the addition of 100 μL, 40 mgL^−1^ sodium carbonate. Chitinase from *Streptomyces griseus* was used as a positive control and a buffer-only condition was used as a negative control. The fluorescence at 450 nm was recorded using a plate reader. Activity was calculated using the equation provided in the product specification sheet. Significance of enzyme activity data was determined using an unpaired two-tailed t-test.

## Acknowledgements

This research was undertaken in part using the MX2 beamline at the Australian Synchrotron, part of ANSTO, and made use of the Australian Cancer Research Foundation (ACRF) detector. This research was undertaken with the assistance of resources from the National Computational Infrastructure (NCI Australia), an NCRIS enabled capability supported by the Australian Government. This work was supported by the Australian Research Council Centre of Excellence in Synthetic Biology (grant CE200100029) and the Australian Research Council Centre for Innovations in Peptide and Protein Science (grant CE200100012).

## Author contributions

S.B.P., J.A.T.D., J.K. and C.J.J. conceived and designed the research. J.A.T.D., O.B.S., J.H. and R.L.F. performed the experiments and data collection. S.B.P. performed the bioinformatic and statistical analyses. S.B.P., R.L.F., J.K. and C.J.J. supervised the research. S.B.P. and C.J.J. wrote the manuscript with input from J.A.T.D. All authors reviewed, edited and approved the final manuscript.

