## Supplementary Data for "The modular evolution of chitinases is governed by coevolution of auxiliary and catalytic domains"

Sacha B. Pulsford, *et al.*

This PDF file includes:

Supplementary Notes 1 to 4

Supplementary Tables 1 to 2

Supplementary Figures 1 to 5

Supplementary Data File Descriptions:

Supplementary File 1: master annotation csv of all GH18 domains

Supplementary File 2: multiple sequence alignment of representative GH18 domains

**Supplementary Note 1: Parsing catalytic domain annotations from InterProScan**

Comprehensive InterProScan annotation reveals a hierarchical organization of GH18-associated signatures, reflecting progressive increases in chitinase complexity from single- to multi-catalytic module sequences. Following the InterProScan workflow, all chitinase domain annotations were deduplicated as described in the Methods section of the main text, preferencing labels PF00704, PS51910, SSF51445 (ranked 1, 2, and 3, respectively); any chitinase domain not associated with one of these top three signatures was assigned the most common signature based on pre-computed global frequencies of the deduped database (Supplementary Table 1). Chitinases with a single catalytic domain (typically in addition to other auxiliary domains) are defined almost exclusively by canonical GH18 catalytic profiles. The dominant signatures are PF00704 (Glycosyl hydrolases family 18), present in 34,021 sequences, and PS51910 (GH18 domain profile), in 3,301 sequences (Supplementary Table 2). These represent the characteristic, conserved catalytic core. Chitinase sequences with two catalytic domains display a marked broadening of GH18 signatures. In addition to continued representation of PF00704 and PS51910, the most frequent and statistically enriched signatures include G3DSA:3.20.20.370, PF06483, SSF51445, and G3DSA:2.10.10.20:FF:000001 (Supplementary Table 3). Additional but less frequent enrichments include GH6-cellulase-like (SSF51989) and glycosidase-related (G3DSA:3.20.20.80) folds (Supplementary Table 3). These combinations indicate that expansion beyond a single GH18 core was primarily achieved through the recruitment of auxiliary catalytic modules, producing bi-modular enzymes that combine chitinase activity with deacetylase- or transglycosidase-like functions.

Three-domain chitinases exhibit the most heterogeneous composition and highest level of functional diversification. They retain enrichment of SSF51445 ((Trans)glycosidases; $q = 9.4 \times 10^{-30}$) but gain multiple distinct chitinase and glycosidase signatures, including G3DSA:2.60.40.10:FF:002602 and G3DSA:2.60.40.10:FF:002609, G3DSA:3.20.20.80, and the canonical PS51910 GH18 profile (Supplementary Table 3). Rare enrichment of PTHR22595 (Chitinase-related), GH6 (SSF51989), and Secreted chitinase profiles further illustrates fusion among multiple hydrolase families (Supplementary Table 3). These recurrent combinations point to tandem duplication and inter-family fusion events that yielded complex, multi-catalytic architectures integrating both canonical and divergent GH18-like modules.

Signatures exclusive to specific classes reinforce this pattern (Supplementary Table 4). Canonical GH18 profiles (PF00704, PS51910) are confined to one-domain enzymes; deacetylase, Chitinase C, and transglycosidase folds typify two-domain sequences; and multiple distinct chitinase subfamilies (e.g., G3DSA:2.60.40.10 variants) appear only in three-domain architectures. Overall, these data support a stepwise evolutionary trajectory of the GH18 family, progressing from single-domain enzymes with a conserved catalytic core to multi-domain chitinases that combine divergent hydrolase and binding modules. The observed pattern suggests that increased domain number correlates with both functional diversification and catalytic complexity, consistent with progressive modular fusion and adaptive broadening of substrate range within the GH18 lineage. Collectively, these patterns support a stepwise evolutionary expansion of domain composition, transitioning from compact GH18 enzymes toward chitinases with multiple catalytic modules integrating divergent chitinase domains that likely enhance substrate range or functional versatility.

**Supplementary Note 2: Sequence domain count details and analyses**

To quantify the distribution of domain counts across all annotated proteins, we first tabulated the number of proteins containing 1, 2, 3, … domains and excluded any zero-domain entries. We modelled the expected frequency of multi-domain architectures using an exponential decay model of the form:

$$P(k) \propto e^{-\lambda(k-1)}\, for\, k \geq 1$$

where $k$ is the number of domains. The decay parameter $\lambda$ was estimated by maximum likelihood from the observed proportion of single-domain proteins relative to the total dataset:

$$\lambda = -ln(\frac{single\ domain\ frequency}{total\ sequences})$$

Expected probabilities were evaluated across the full observed domain range, normalized to sum to one, and multiplied by the total number of proteins to derive the expected domain-count frequencies under this exponential model. To test whether the empirical data deviated significantly from this expectation, we performed a chi-squared goodness-of-fit test using observed versus expected frequencies for multi-domain architectures (domain count ≥2), with the expected values rescaled to match the total number of multi-domain proteins ($\chi^2$ = 18891.8, p = 0.00e+00). To specifically assess enrichment of three-domain architectures relative to two-domain architectures, we performed a one-sided binomial test in which the number of three-domain proteins was compared to the combined pool of two- and three-domain proteins under a null probability of 0.5 (odds ratio = 0.97, p = 9.92e-01). Finally, we calculated standardized Pearson residuals for each domain count class as:

$$(Observed - Expected)/\sqrt{Expected}$$

to identify domain classes that contributed most strongly to deviations from the exponential expectation (Supplementary Table 6). The most significant residual was that of three-domain sequences (z = 120.9).

### Supplementary Note 3: DVI analysis summary

*Domain adjacency and GH18-specific promiscuity metric.* Domain architectures were derived from InterProScan cleaned annotation outputs (as described above). For each non-GH18 domain, we quantified two properties: (i) how often it co-occurred with GH18 in a protein, and (ii) how many distinct domain neighbours it exhibited within GH18-containing architectures. To count co-occurrence, we first constructed, for each GH18-positive protein, the set of unique non-GH18 domains present in its architecture. For each domain *d* in this set, we incremented a co-occurrence count $n_{(\mathrm{GH18,\ d})}$ by one. This ensures that $n_{(\mathrm{GH18,\ d})}$ reflects the number of GH18-containing proteins in which *d* appears at least once, independent of its multiplicity within a single architecture.

To quantify neighbour diversity, we treated the ordered domain list of each GH18-containing protein as a linear architecture and recorded, for every non-GH18 domain *d* in that list, its immediate left and right neighbours (where present). For each appearance of *d*, the domains at positions $i - 1$ and $i + 1$ were considered candidate neighbours; if a neighbour existed and was not identical to *d*, it was added to the neighbour set of *d*. Across all proteins, this yielded, for each domain *d*, a set of distinct neighbours, whose cardinality defined the total neighbour diversity $T_{(\mathrm{GH18,\ d})}$ (i.e. the number of unique domains observed adjacent to *d* within GH18-containing architectures). To convert these counts into relative measures, the neighbour diversity of each domain *d* was normalised by the total neighbour diversity summed across all domains in the dataset. Specifically, we defined:

$$\pi_{(GH18,d)} = T_{(GH18,d)} \;/\; \Sigma_{\mathrm{j}}\, T_{(GH18,j)}$$

where indexes all non-GH18 domains considered in the analysis (i.e., the same domain set to which *d* belongs). In this formulation, *d* refers to the focal domain for which the normalised diversity is being calculated, whereas *j* ranges over every domain in the dataset to provide the denominator representing the total neighbour diversity. In parallel, the co-occurrence frequency of each domain was normalised by the total number of GH18-containing proteins ($f_{(\mathrm{GH18},\ d)}$), reflecting how common *d* is among GH18 architectures.

To obtain a GH18-specific promiscuity score (φ), we adapted the Domain Versatility Index framework by comparing the neighbour diversity of a domain to its abundance. For each domain *d*, we computed:

$$\varphi_{(GH18,d)} = \pi_{(GH18,d)} \cdot ln\left(\pi_{(GH18,d)} \;/\; f_{(GH18,d)}\right)$$

setting $\varphi_{(GH18,\ d)} = 0$ when either $\pi_{(GH18,\ d)}$ or $f_{(GH18,\ d)}$ was zero. This measure assigns higher scores to domains that, relative to how frequently they appear in GH18-containing proteins, are associated with a disproportionately diverse set of neighbouring domains, and therefore are more likely to participate in a wide range of GH18 architectures. Finally, for each domain we recorded: the domain identifier, its GH18 co-occurrence count ($n_{(GH18,\ d)}$), its neighbour diversity ($T_{(GH18,\ d)}$), its normalised neighbour contribution ($\pi_{(GH18,\ d)}$), and its GH18-specific promiscuity score ($\varphi_{(GH18,\ d)}$).

*Statistical enrichment of promiscuous domains across SSN clusters and biological categories.* To evaluate whether particular functional categories were disproportionately represented among GH18-associated domains, we performed two complementary enrichment analyses. First, to assess enrichment of functional classes among domains identified as promiscuous partners or housekeeper partners of GH18 catalytic domains, we compared their category frequencies against those observed across the full annotated domain set (denoted 'Global'). For each category, we tabulated (i) the number of promiscuous/housekeeper domains assigned to that category and (ii) the number of domains annotated with the same category across the global dataset. These counts formed the basis of a 2×2 contingency table contrasting the representation of each category. Statistical enrichment was tested using Fisher's exact test (one-sided, alternative = "greater"), and *p*-values were corrected for multiple testing using the Benjamini–Hochberg false discovery rate (FDR). An enrichment ratio, calculated as the ratio of category counts between the promiscuous/housekeeper and background sets (with a pseudocount of +1), was used to provide an effect-size estimate. An equivalent calculation was then made comparing the promiscuous and housekeeper domain datasets. All analyses were performed in Python using SciPy and statsmodels.

### Supplementary Note 4: Rooting the GH18 chitinase phylogeny in the absence of a clear outgroup

For the GH18 chitinase dataset analysed here, no clear outgroup is available: all sequences are bacterial or archaeal, and archaeal sequences do not form a monophyletic cluster but instead occur across multiple distal clades. This distribution precludes confident designation of any single lineage as a reference for polarity and motivates the use of internal rooting criteria. The phylogeny shown in the main text is therefore presented midpoint-rooted to provide a consistent orientation of the ML topology.

To evaluate the stability of this orientation, we inferred five independent ML phylogenies using IQ-TREE3 from the same multiple sequence alignment (Methods) and compared (i) midpoint rooting to (ii) Minimal Ancestral Deviation (MAD) rooting, which places the root to minimize global deviation from broadly clock-like behaviour across the tree, and (iii) rooting under the nonreversible nQ.pfam substitution model, under which the root becomes statistically identifiable from the sequence data because likelihood depends on substitution direction along branches (Supplementary Figure 3 and Supplementary Figure 4). Across the five replicate searches, the unrooted backbone topology was highly consistent, indicating that major clade structure and inter-clade relationships are not contingent on a single tree-search trajectory.

MAD-rooted topologies of all independent replicate trees were broadly concordant with the corresponding midpoint-rooted topologies and recovered the same key groupings defined by InDel Classes (Supplementary Fig. 3). In particular, InDel Classes A and C–G consistently formed monophyletic groups, as observed under midpoint rooting. By contrast, InDel Class B was consistently positioned at or near the inferred root and was typically polyphyletic, appearing as staggered basal sequences rather than a single cohesive clade. Despite this, the internal ordering of subclades within the broader InDel Class B assemblage was consistent in 4/5 replicate topologies: InDel Class B sequences comprised the most basal groups, transitioning through InDel Class A, with subsequent radiation to InDel Classes C–G. A single replicate (replicate 1) deviated modestly from this dominant pattern, with a subset of InDel Class B sequences forming a tight basal cluster at the root while additional InDel Class B sequences also appeared as a clade between InDel Class A and the C–G assemblage. Importantly, this deviation did not disrupt the overarching agreement between midpoint and MAD orientations, supporting the interpretation that the apparent "centre" of the tree is not being driven by an idiosyncratic long branch; rather, multiple rate-based criteria favour the same broad polarity.

Rooting under nQ.pfam produced a more heterogeneous set of rooted topologies across replicates; no single fully dominant rooted arrangement was observed (Supplementary Figure 4). Nevertheless, the results were strongly consistent at the level most relevant to biological interpretation: InDel Classes remained largely intact and retained the same overarching relationships seen in midpoint/MAD-rooted trees, with InDel Classes A and B predominantly occupying basal positions and other InDel Classes radiating outward from these inferred ancestral sequence groups. Two replicates placed a tight cluster of InDel Class A at the root and recovered two prominent monophyletic groupings, with InDel Classes C–E forming one clade and InDel Classes F–G forming another. Replicate 4 closely resembled the dominant MAD topology, placing InDel Class B at the base, whereas replicate 1 more closely resembled the midpoint-rooted orientations. The remaining replicate (replicate 3) differed primarily in the placement of InDel Class C, with sequences typically assigned to Class C–E occupying a more basal position than in the other nQ.pfam runs. Taken together, the nQ.pfam analyses reinforce that, even when rooted topologies vary in fine detail, the principal signal (basal enrichment of InDel Classes A/B and stable higher-level grouping of C–G) remains consistent with midpoint and MAD rooting.

Collectively, agreement among midpoint rooting (a geometric heuristic), MAD (a global rate-regularity criterion), and nQ.pfam (a likelihood-based, non-reversible rooting framework) supports the use of the midpoint-rooted representation as a clear primary visualization and indicates that the core conclusions that depend on major clade structure and InDel-defined grouping are robust to alternative rooting strategies. Nevertheless, because no external outgroup is available, explicit statements requiring evolutionary polarity (for example, designation of "ancestral" versus "derived" clades or the temporal ordering of trait acquisition) are presented as root-contingent hypotheses supported by concordant internal criteria rather than as outgroup-anchored certainty.

**Supplementary Table 1.** Top ten most frequent chitinase-related signature counts across the dataset prior to label deduplication.

| Signature | Count |
|---|---|
| G3DSA:3.20.20.80 | 40713 |
| SSF51445 | 40009 |
| PS51910 | 39036 |
| PF00704 | 36361 |
| SM00636 | 31254 |
| G3DSA:3.10.50.10 | 23653 |
| PTHR11177 | 16418 |
| PIRSR001103-1 | 13287 |
| SSF54556 | 11782 |
| cd06548 | 11059 |

**Supplementary Table 2.** Counts of sequences with 1, 2 or 3 catalytic domains (GH18 domains) across all GH18-related InterProScan signatures after label deduplication.

| | | GH18 domains | | |
|---|---|---|---|---|
| **Signature** | **Description** | 1 | 2 | 3 |
| G3DSA:2.10.10.20:FF:000001 | Secreted chitinase | 0 | 129 | 4 |
| G3DSA:2.60.40.10:FF:002602 | Chitinase | 0 | 4 | 7 |
| G3DSA:2.60.40.10:FF:002609 | Chitinase | 0 | 0 | 7 |
| G3DSA:2.60.40.10:FF:003164 | Chitinase A | 0 | 1 | 0 |
| G3DSA:3.10.50.10 | — | 0 | 4 | 0 |
| G3DSA:3.20.20.370 | Glycoside hydrolase/deacetylase | 0 | 1389 | 3 |
| G3DSA:3.20.20.80 | Glycosidases | 0 | 5 | 4 |
| G3DSA:3.30.1490.230 | — | 0 | 1 | 0 |
| G3DSA:3.30.20.10:FF:000003 | Chitinase | 0 | 2 | 0 |
| PF00704 | Glycosyl hydrolases family 18 | 34021 | 2234 | 97 |
| PF06483 | Chitinase C | 0 | 231 | 0 |
| PIRSR001103-1 | — | 0 | 12 | 0 |
| PS51910 | Glycosyl hydrolases family 18 (GH18) domain profile | 3301 | 176 | 23 |
| PTHR10963 | Glycosyl hydrolase related | 0 | 5 | 0 |
| PTHR22595 | Chitinase-related | 0 | 4 | 1 |
| SSF51445 | (Trans)glycosidases | 1 | 187 | 36 |
| SSF51989 | Glycosyl hydrolases family 6, cellulases | 0 | 10 | 1 |

**Supplementary Table 3.** Signatures enriched within multi-domain chitinases. Fisher's exact tests were performed to assess enrichment of individual InterProScan signatures within proteins containing two or three total domains. Odds ratios reflect the ratio of odds of a signature occurring within the tested class versus outside it; q-values are FDR-corrected p-values. Proportions denote the fraction of sequences in-class and out-of-class containing each signature. Only signatures significant at FDR < 0.05 are shown. No significant enrichments were detected for single-domain chitinases; only Class 2 (2 catalytic domains) and Class 3 (3 catalytic domains) are shown.

| Domain class | Enriched signature accession | Signature description | Odds ratio | q-value (FDR) | Proportion in class | Proportion outside | Δ proportion | Proteins in class | Proteins outside |
|---|---|---|---|---|---|---|---|---|---|
| **Class 2** | G3DSA:3.20.20.370 | Glycoside hydrolase/deacetylase | $3.2 \times 10^{4}$ | 0 | 0.632 | $5 \times 10^{-5}$ | +0.632 | 1 389 | 2 |
| | PF06483 | Chitinase C | ∞ | $6.2 \times 10^{-295}$ | 0.105 | 0 | +0.105 | 231 | 0 |
| | SSF51445 | (Trans)glycosidases | 165 | $5.0 \times 10^{-210}$ | 0.085 | $5.6 \times 10^{-4}$ | +0.085 | 187 | 21 |
| | G3DSA:2.10.10.20:FF:000001 | Secreted chitinase | $1.2 \times 10^{3}$ | $1.0 \times 10^{-159}$ | 0.059 | $5 \times 10^{-5}$ | +0.059 | 129 | 2 |
| | PIRSR001103-1 | — | ∞ | $3.0 \times 10^{-15}$ | 0.005 | 0 | +0.005 | 12 | 0 |
| | SSF51989 | GH6 cellulases | 171 | $8.6 \times 10^{-12}$ | 0.005 | $3 \times 10^{-5}$ | +0.005 | 10 | 1 |
| | PTHR10963 | Glycosyl hydrolase-related | ∞ | $1.4 \times 10^{-6}$ | 0.002 | 0 | +0.002 | 5 | 0 |
| | G3DSA:3.10.50.10 | — | ∞ | $2.1 \times 10^{-5}$ | 0.002 | 0 | +0.002 | 4 | 0 |
| | PTHR22595 | Chitinase-related | 68 | $9.1 \times 10^{-5}$ | 0.002 | $3 \times 10^{-5}$ | +0.002 | 4 | 1 |
| | G3DSA:3.20.20.80 | Glycosidases | 21 | $9.9 \times 10^{-5}$ | 0.002 | $1 \times 10^{-4}$ | +0.002 | 5 | 4 |
| **Class 3** | SSF51445 | (Trans)glycosidases | 102 | $9.4 \times 10^{-30}$ | 0.328 | 0.0048 | +0.323 | 20 | 188 |
| | G3DSA:2.60.40.10:FF:002609 | Chitinase | ∞ | $1.3 \times 10^{-19}$ | 0.115 | 0 | +0.115 | 7 | 0 |

| | | | | | | | | | |
|---|---|---|---|---|---|---|---|---|---|
| | G3DSA:2.60.40.10: FF:002602 | Chitinase | $1.3 \times 10^{3}$ | $2.8 \times 10^{-17}$ | 0.115 | $1 \times 10^{-4}$ | +0.115 | 7 | 4 |
| | G3DSA:3.20.20.80 | Glycosidases | 555 | $2.9 \times 10^{-9}$ | 0.066 | $1 \times 10^{-4}$ | +0.065 | 4 | 5 |
| | PS51910 | GH18 domain profile | 5.85 | $1.6 \times 10^{-8}$ | 0.361 | 0.088 | +0.273 | 22 | 3 476 |
| | PTHR22595 | Chitinase-related | 165 | $2.3 \times 10^{-2}$ | 0.016 | $1 \times 10^{-4}$ | +0.016 | 1 | 4 |
| | G3DSA:2.10.10.20: FF:000001 | Secreted chitinase | 10.4 | $3.9 \times 10^{-2}$ | 0.033 | 0.003 | +0.030 | 2 | 129 |
| | SSF51989 | GH6 cellulases | 65.9 | $3.9 \times 10^{-2}$ | 0.016 | 0.0003 | +0.016 | 1 | 10 |

**Supplementary Table 4**. Signatures exclusive to single GH18 domain sequences (Class 1), double GH18 domain sequences (Class 2) or triple GH18 domain sequences (Class 3).

| **Signature** | **Class 1** | **Class 2** | **Class 3** |
|---|---|---|---|
| PF06483 (Chitinase C) | 0 | 231 | 0 |
| PIRSR001103-1 | 0 | 12 | 0 |
| PTHR10963 (GH-related) | 0 | 5 | 0 |
| G3DSA:3.10.50.10 | 0 | 4 | 0 |
| G3DSA:3.30.20.10:FF:000003 | 0 | 2 | 0 |
| cd00413 (GH16) | 0 | 2 | 0 |
| G3DSA:2.60.40.10:FF:003164 | 0 | 1 | 0 |
| G3DSA:3.30.1490.230 | 0 | 1 | 0 |
| G3DSA:2.60.40.10:FF:002609 | 0 | 0 | 7 |

**Supplementary Table 5**. Top 15 most frequent auxiliary domain signature count across the dataset after de-duplication. Complete signature details available as supplementary datafile.

| Signature | Count |
|---|---|
| PF02839 | 5928 |
| PF01476 | 5828 |
| PF00041 | 5527 |
| PF00553 | 3375 |
| PS51257 | 2733 |
| PF17957 | 2686 |
| SM00495 | 2650 |
| PF02018 | 2611 |
| G3DSA:2.60.40.10 | 2188 |
| PF00395 | 1953 |
| PS51318 | 1657 |
| PF07833 | 1226 |
| PF08239 | 914 |
| PF08329 | 909 |
| PF00535 | 719 |

**Supplementary Table 6.** The number of domains per sequence in the cleaned GH18 dataset. The 'Observed counts' column refers to the actual number of sequences observed to comprise a specific number of domains. The 'Expected counts' column refers to the corresponding expected frequency under exponential decay. Residuals quantify the extent of the deviation between observed and expected counts. To confirm sequences with extreme domain numbers (>8 domains in a sequence), manual analysis was conducted by inferring the structure of arbitrarily selected sequences in each of these classes with Alphafold3 to confirm the existence of such extensive architecture. These sequences indeed possess a remarkable number of auxiliary domains, in the most extreme case, A0A179SQD3 comprises 9 LysM domains, 7 immunoglobulin repeats, a Ca2+ tandem repeat and a single GH18 domain. It is unlikely these extensive sequences exist as bead-on-a-string-like assembly *in vivo*, rather assembling into more complex oligomers. Indeed, Ca2+ motif is associated with inducing rigidity in large adhesion-like assemblies of multiple immunoglobulin-like domains in the biofilm-associated protein Bap of *Acinetobacter baumannii* (which can exceed 8000 amino acids in length), the calcium-stabilized ice-binding adhesin of the Antarctic bacterium *Marinomonas primoryensis*, and the giant calcium-binding adhesin SiiE of *Salmonella enterica*. Considering this, it may be that higher order assemblies coordinated by such extensive architecture exist as a general mechanism for binding substrates.

| Domain number | Observed counts | Expected counts | Residual |
|---|---|---|---|
| 1 | 13207 | 26375.00007 | -81.081838 |
| 2 | 9819 | 8800.329087 | 10.858865 |
| 3 | 9486 | 2936.33334 | 120.869443 |
| 4 | 4405 | 979.742166 | 109.430244 |
| 5 | 1238 | 326.902501 | 50.39132 |
| 6 | 652 | 109.074866 | 51.984947 |
| 7 | 498 | 36.394112 | 76.51662 |
| 8 | 167 | 12.143324 | 44.43868 |
| 9 | 36 | 4.051763 | 15.871753 |
| 10 | 26 | 1.351918 | 21.198635 |
| 11 | 15 | 0.451083 | 21.662182 |
| 12 | 17 | 0.150509 | 43.431528 |
| 13 | 10 | 0.050219 | 44.399555 |
| 14 | 3 | 0.016756 | 23.046269 |
| 15 | 1 | 0.005591 | 13.299146 |
| 16 | 1 | 0.001865 | 23.109711 |
| 17 | 0 | 0.000622 | -0.024949 |
| 18 | 1 | 0.000208 | 69.375926 |

**Supplementary Table 7.** All auxiliary domains were manually assigned to broad biological categories for downstream analysis based on signature annotation information available on the InterPro database.

| Functional class | Functional category | Description | Number of unique auxiliary domain signatures associated with category |
|---|---|---|---|
| Substrate-recognition and targeting modules | Carbohydrate-binding modules (CBMs) | Glycan-binding domains that tether enzymes to chitin or related polysaccharides, increasing local substrate concentration and enhancing processivity. | 36 |
| | Lectins (β-trefoil / jacalin / C-type / others) | Sugar-recognition modules that bind specific glycan motifs, often contributing to substrate specificity, adhesion, or recognition of extracellular carbohydrate structures. | 18 |
| | Peptidoglycan-/cell-wall binding and SH3b group | Domains that recognise peptidoglycan or cell-wall components, supporting cell-envelope targeting or stable cell-surface association. | 27 |
| Structural spacers, scaffolds and architectural supports | Ig-like & related β-sandwich scaffolds | β-sandwich modules (e.g., Fn3, Ig-like, PKD) that act as structural spacers, rigidity elements, or protein–protein interaction surfaces within multidomain enzymes. | 70 |
| | Repeat/stalk & coiled-coil/IDRs | Repetitive motifs, coiled-coil elements, and intrinsically disordered regions that function as flexible or extended spacers positioning catalytic domains relative to their substrates. | 22 |
| Catalytic and biochemical partner systems | Catalytic partner domains (non-GH18) | Auxiliary enzymatic modules, including glycosyltransferases and NDP-sugar–modifying enzymes, that provide additional catalytic chemistry adjacent to the GH18 core. | 70 |
| | Redox/stress/metal helpers | Domains associated with redox regulation, oxidative stress response, or metal | 10 |

| | | | |
|---|---|---|---|
| | | binding, potentially stabilising extracellular enzymes or modulating catalytic activity. | |
| | Cellulosome / cellulolytic apparatus add-ons | Accessory modules typical of cellulosomal systems that contribute to polysaccharide-degrading complex assembly or scaffold interactions. | 3 |
| Secretion, localisation and cell-envelope association | Surface anchoring & secretion/sorting systems | Motifs and domains involved in secretion, membrane anchoring, or cell-surface localisation, including Tat signal peptides and lipoprotein attachment sites. | 24 |
| | Outer-layer/periplasmic envelope helpers | Periplasmic or envelope-associated modules implicated in surface maintenance, periplasmic trafficking, or folding of secreted enzymes. | 6 |
| Regulatory and rare fusion elements | DNA/RNA-binding & regulators occasionally fused | Infrequent nucleic acid-binding or regulatory domains that appear as rare domain fusions, potentially adding transcriptional, localisation, or regulatory layers to GH18-associated proteins. | 7 |
| Poorly Characterised or Unannotated Elements | Unknowns / DUFs / broad structure-only calls | Uncharacterised domains or structure-only predictions for which no clear biochemical or cellular function can yet be assigned. | 72 |

**Supplementary Table 8.** Details of auxiliary domains enrichment in the promiscuous dataset relative to the global dataset. No broad functional category remained significantly enriched among promiscuous domains after correction for multiple testing. However, several categories showed directional trends (odds ratios > 1). Ig-like β-sandwich scaffolds exhibited the strongest tendency toward enrichment, with lectins and carbohydrate-binding modules (CBMs) also displaying moderate positive biases. Conversely, categories such as unknown/DUF-like domains and surface-anchoring/secretion systems appeared numerically under-represented (odds ratio < 1), although statistical support for depletion was weak.

| Broad category | Promiscuous count | Global count | Odds ratio |
|---|---|---|---|
| Ig-like & related β-sandwich scaffolds (adhesion, spacing) | 14 (35%) | 70 (19.2%) | 2.27 |
| Lectins (β-trefoil / jacalin / C-type / others) | 4 (10.0%) | 18 (4.9%) | 2.14 |
| Carbohydrate-binding modules (CBMs) — glycan grips | 7 (17.5%) | 36 (9.9%) | 1.94 |
| Repeat/stalk & coiled-coil/IDRs (mechanical spacing) | 2 (5%) | 22 (6.0%) | 0.82 |
| Catalytic partner domains (non-GH18) | 6 (15%) | 70 (19.2%) | 0.74 |
| Peptidoglycan-/cell-wall binding and SH3b group | 2 (5.0%) | 27 (7.4%) | 0.66 |
| Unknowns / DUFs / broad structure-only calls | 4 (10.0%) | 72 (19.7%) | 0.45 |
| Surface anchoring & secretion/sorting systems | 1 (2.5%) | 24 (6.6%) | 0.36 |
| Redox/stress/metal helpers | 0 (0%) | 10 (2.7%) | 0.00 |
| DNA/RNA-binding & regulators occasionally fused | 0 (0%) | 7 (1.9%) | 0.00 |
| Outer-layer/periplasmic envelope helpers | 0 (0%) | 6 (1.6%) | 0.00 |
| Cellulosome / cellulolytic apparatus add-ons | 0 (0%) | 3 (0.8%) | 0.00 |

**Supplementary Table 9.** Details of auxiliary domains enrichment in the housekeeper dataset relative to the global dataset. No broad functional category showed significant enrichment or depletion among housekeeper-associated domains after correction for multiple testing. Nevertheless, several categories displayed directional trends: CBMs were more frequent in the housekeeper set relative to the global domain background (odds ratio > 1), with surface anchoring/sorting systems and peptidoglycan-/cell-wall binding SH3b domains also exhibiting moderate positive biases. Conversely, uncharacterized/DUF-like domains appeared under-represented among housekeepers (odds ratios < 1), although statistical support for depletion was weak.

| Broad category | Housekeeper count | Global count | Odds ratio |
|---|---|---|---|
| Carbohydrate-binding modules (CBMs) — glycan grips | 5 (23.8%) | 36 (9.9%) | 2.856 |
| Peptidoglycan-/cell-wall binding and SH3b group | 3 (14.3%) | 27 (7.4%) | 2.086 |
| Surface anchoring & secretion/sorting systems | 3 (14.3%) | 24 (6.6%) | 2.368 |
| Catalytic partner domains (non-GH18) | 3 (14.3%) | 70 (19.2%) | 0.702 |
| Cellulosome / cellulolytic apparatus add-ons | 0 (0.00%) | 3 (0.8%) | 0.000 |
| DNA/RNA-binding & regulators occasionally fused | 0 (0.00%) | 7 (1.9%) | 0.000 |
| Ig-like & related β-sandwich scaffolds (adhesion, spacing) | 4 (19.1%) | 70 (19.2%) | 0.992 |
| Lectins (β-trefoil / jacalin / C-type / others) | 1 (4.8%) | 18 (4.9%) | 0.964 |
| Outer-layer/periplasmic envelope helpers | 0 (0.00%) | 6 (1.6%) | 0.000 |
| Redox/stress/metal helpers | 0 (0.00%) | 10 (2.7%) | 0.000 |
| Repeat/stalk & coiled-coil/IDRs (mechanical spacing) | 0 (0.00%) | 22 (6.0%) | 0.000 |
| Unknowns / DUFs / broad structure-only calls | 2 | 72 (19.7%) | 0.428 |

**Supplementary Table 10.** Promiscuous vs housekeeper domain enrichment patterns. Although limited sample sizes preclude firm statistical conclusions when comparing housekeeper and promiscuous datasets broad trends pertain to directional trends. Promiscuous GH18 domains are enriched in Ig-like spacers, repeats, and lectins domains that tolerate recombination and facilitate new architectures. Comparatively, housekeepers retain the surface anchoring/sorting systems, peptidoglycan-/SH3b envelope anchors, and CBM-linked organization.

| Broad Category | Enrichment Pattern (promiscuous vs housekeeper) | Odds ratio (promiscuous vs housekeeper) | Interpretation |
|---|---|---|---|
| **Ig-like & related β-sandwich scaffolds (adhesion, spacing)** | Higher in promiscuous (35 % vs 19 %) | 2.29 | Over-represented in promiscuous architectures. They act as flexible connectors, mechanically decoupling catalytic and binding modules, ideal for recombination and domain shuffling. |
| **Lectins (β-trefoil / jacalin / C-type / others)** | Mildly higher in promiscuous (10 % vs 5 %) | 2.22 | Over-represented in promiscuous architectures. Grafting to catalytic cores may broaden target recognition, promoting secondary binding specificity diversification. |
| **Carbohydrate-binding modules (CBMs)** | Slightly lower (17.5 % vs 24 %) | 0.68 | CBMs are common in both, but relatively depleted in promiscuous cases, consistent with them being ancient, conserved “housekeeping” attachments rather than drivers of modular novelty. |
| **Peptidoglycan / SH3b group** | Lower (5 % vs 14 %) | 0.32 | Enrichment in housekeepers implies cell-wall localization is a stable, non-recombined trait. |
| **Surface anchoring & secretion systems** | Much lower (2.5 % vs 14 %) | 0.15 | Enrichment of housekeepers indicates membrane-anchoring is a stable, non-recombined trait. |
| **Catalytic partner domains (non-GH18)** | Equal (~15 %) | 1.06 | Both classes maintain a similar background level of accessory catalytic domains. |
| **Repeat / coiled-coil / IDR domains** | Present only in promiscuous | ∞ | Intrinsically disordered or repeat regions appear exclusively in promiscuous set (though n = 2). Such regions promote flexibility and fusion tolerance, consistent with recombination hotspots. |
| **Unknown / DUFs** | Equal (~10%) | 1.06 | No difference. |

**Supplementary Table 11.** Phylogeny replicate information and statistics. Clade labels correspond to those clades annotated on the tree in the main text in Figure 3a. All replicates were independently inferred from the same multiple sequence alignment using IQTREE3.

| Tree | Log (Likelihood) | ultrafast Bootstraps values for key clades | p-AU |
|---|---|---|---|
| 1 | -322956 | Basal 1 (clades I and II): 96<br>Basal 2 (clades III-V): 25<br>Basal 3 (clades IV and V): 100<br>Clade I:100<br>Clade II:97<br>Clade III:55<br>Clade IV:97<br>Clade V:97 | 0.494 |
| 2 | -322960 | Basal 1 (clades I and II):63<br>Basal 2 (clades III-V): 99<br>Basal 3 (clades IV and V): 100<br>Clade I: 100<br>Clade II: 63<br>Clade III: 83<br>Clade IV: 96<br>Clade V: 97 | 0.442 |
| 3 | -322970 | Basal 1 (clades I and II): 58<br>Basal 2 (clades III-V): 99<br>Basal 3 (clades IV and V): 100<br>Clade I: 100<br>Clade II: 62<br>Clade III: 81<br>Clade IV:96<br>Clade V: 92 | 0.365 |
| 4 | -322955 | Basal 1 (clades I and II):31<br>Basal 2 (clades III-V):100<br>Basal 3 (clades IV and V): 100<br>Clade I:100<br>Clade II:35<br>Clade III:76<br>Clade IV: 95<br>Clade V: 95 | 0.527 |
| 5 | -322954 | Basal 1 (clades I and II): 59<br>Basal 2 (clades III-V): 98<br>Basal 3 (clades IV and V): 100<br>Clade I: 100<br>Clade II: 62<br>Clade III: 78<br>Clade IV: 96<br>Clade V: 88 | 0.528 |

**Supplementary Table 12.** Crystallography data collection and refinement statistics for the three chitinase domain structures reported here.

| **Structure** | EC17625 | SV2589 | Vs05695 |
|---|---|---|---|
| **PDB ID** | 9BUG | 9BUF | 9OJL |
| **Data collection** | | | |
| Space group | $P\ 2_1\ 2_1\ 2$ | I 1 2 1 | $P\ 2_1\ 2_1\ 2_1$ |
| Cell dimensions | | | |
| *a, b, c (Å)* | 83.06 90.45 54.58 | 77.53 54.77 144.42 | 109.92 154.03 156.60 |
| α, β, γ (°) | 90.00 90.00 90.00 | 90.00 92.13 90.00 | 90.00 90.00 90.00 |
| Resolution (*Å*) | 46.73 - 1.30 (1.32 - 1.30)* | 41.57 - 1.75 (1.78 - 1.75) | 49.44 - 1.80 (1.83 - 1.80) |
| Rmerge | 0.083 (2.258) | 0.244 (2.302) | 0.201 (2.092) |
| Rpim | 0.024 (0.623) | 0.101 (0.929) | 0.056 (0.582) |
| I/σI | 14.4 (1.2) | 6.2 (1.0) | 10.7 (1.3) |
| $CC_{1/2}$ | 0.999 (0.652) | 0.993 (0.504) | 0.998 (0.358) |
| Completeness (%) | 97.9 (96.0) | 99.9 (100.0) | 100.0 (100.0) |
| Redundancy | 13.5 (13.9) | 6.8 (7.0) | 13.9 (13.8) |
| **Refinement** | | | |
| Resolution (*Å*) | 46.73 - 1.30 (1.31 - 1.30) | 41.87 - 1.75 (1.78 - 1.75) | 49.44 - 1.80 (1.82 - 1.80) |
| No. reflections | 99316 (9631) | 61231 (6087) | 244980 (24273) |
| $R_{work}/R_{free}$ | 0.157/0.184 (0.324/0.351) | 0.195/0.232 (0.297/0.348) | 0.168/0.209 (0.325/0.35) |
| **No. atoms** | | | |
| Protein | 3242 | 4334 | 15176 |
| Ligand/ion | 38 | 14 | 122 |
| Water | 542 | 545 | 2400 |
| **B-factors (overall)** | 24.08 | 22.08 | 30.10 |
| Protein | 22.03 | 21.36 | 28.86 |
| Ligand/ion | 57.31 | 34.84 | 39.20 |
| Water | 34.02 | 27.54 | 37.50 |
| **R.m.s. deviations** | | | |
| Bond lengths (*Å*) | 0.006 | 0.007 | 0.013 |
| Bond angles (°) | 0.944 | 0.884 | 1.119 |

*Values in parentheses are for the highest resolution shell

**Supplementary Table 13**. Chitinases cloned and heterologously expressed in *E. coli* BL21 (DE3) in this study. Listed are each construct, its assigned clade, whether soluble protein could be purified, and whether the construct was carried forward for kinetic characterization.

| Uniprot ID | Clade | Molecular Weight (kDa) | Purified | Kinetic Analysis |
|---|---|---|---|---|
| A0A420CKH5 | I | 34 | No | No |
| A0A5M7C9P2 | I | 35 | Yes | Yes |
| F2R3N3 | I | 35 | Yes | Yes |
| F2R3N3_FL | I | 51 | Yes | Yes |
| A0A1G7R0D1_2 | II | 40 | Yes | Yes |
| A0A505D7J4 | II | 34 | Yes | Yes |
| A0A837NXJ2 | II | 39 | Yes | Yes |
| A0A376E139 | III | 52 | No | No |
| A0A8E1YAV4 | III | 50 | No | No |
| B7VRF8 | III | 58 | No | No |
| A0A385THE8 | III | 49 | No | No |
| A0A0R0B9V6 | III | 47 | Yes | Yes |
| A0A429UIK5 | III | 44 | No | No |
| A0A8J3RIB1 | III | 50 | Yes | Yes |
| A0A855LX82 | III | 48 | Yes | Yes |
| A0A2G6TAK0 | III | 42 | Yes | Yes |
| A0A1Y6KW18 | III | 57 | No | No |
| H8MUM6 | III | 42 | Yes | Yes |
| A0A1G7R0D1_1 | III | 53 | Yes | No |
| A0A0R0B9V6_FL | III | 72 | Yes | Yes |
| A0A429UIK5_FL | III | 64 | No | No |
| A0A429UIK5_CBM2 | III | 52 | No | No |
| A0A7H8PJW8 | IV | 48 | Yes | Yes |
| A0A1Y2RHE5 | IV | 38 | No | No |
| A0A7Z7CEU3 | V | 35 | No | No |
| A0A2N9Z3X8 | V | 39 | No | No |
| A0A5C1HUI1 | V | 40 | No | No |
| A0A4Z1AH30 | V | 53 | No | No |
| A0A418T651 | V | 41 | No | No |
| A0A1G7R0D1_FL | II&III | 144 | Yes | No |

**Supplementary Table 14.** Most abundant phyla contributing to each catalytic InDel class. For each InDel class (A to G), the five most common contributing phyla are shown with their raw counts (number of sequences assigned to that phylum within the class) and corresponding proportion of the total sequences in that class (%). This summary highlights major shifts in phylum-level composition across InDel classes.

| InDel Class | Phylum | count | Proportion of InDel Class (%) |
|---|---|---|---|
| **A** | Actinomycetota | 2937 | 86.00 |
| | Pseudomonadota | 257 | 7.50 |
| | Bacillota | 108 | 3.20 |
| | Methanobacteriota | 55 | 1.60 |
| | Chloroflexota | 26 | 0.76 |
| **B** | Actinomycetota | 4148 | 53.00 |
| | Pseudomonadota | 1483 | 19.00 |
| | Bacillota | 906 | 12.00 |
| | Bacteroidota | 876 | 11.00 |
| | Chloroflexota | 89 | 1.10 |
| **C** | Pseudomonadota | 4104 | 39.00 |
| | Bacillota | 2091 | 20.00 |
| | Bacteroidota | 2067 | 20.00 |
| | Actinomycetota | 938 | 8.90 |
| | NaN | 192 | 1.80 |
| **D** | Actinomycetota | 1760 | 75.00 |
| | Pseudomonadota | 411 | 18.00 |
| | Myxococcota | 54 | 2.30 |
| | Acidobacteriota | 49 | 2.10 |
| | Chloroflexota | 29 | 1.20 |
| **E** | Actinomycetota | 2342 | 98.00 |
| | Pseudomonadota | 33 | 1.40 |
| | Myxococcota | 5 | 0.21 |
| | Chloroflexota | 4 | 0.17 |
| **F** | Bacteroidota | 352 | 29.00 |
| | Chloroflexota | 160 | 13.00 |
| | NaN | 92 | 7.60 |
| | Fidelibacteriota | 65 | 5.40 |
| | Ignavibacteriota | 59 | 4.90 |
| **G** | Bacillota | 7469 | 66.00 |
| | Actinomycetota | 834 | 7.40 |
| | Pseudomonadota | 637 | 5.70 |
| | Bacteroidota | 560 | 5.00 |
| | Spirochaetota | 389 | 3.50 |

**Supplementary Table 15.** Summary of phyla that are most strongly enriched or depleted in each catalytic InDel class (A to G). For each InDel–phylum combination, Count reports the number of sequences assigned to that phylum in the given InDel class, with the percentage of all sequences in that class shown in brackets. $Log_2$(odds ratio) values quantify over- or under-representation of each phylum in that class relative to all other classes combined, based on 2×2 contingency tables with a continuity correction. p values are from Fisher's exact test, and FDR (p-value) values are Benjamini–Hochberg–adjusted false discovery rates used to identify significantly enriched (positive $log_2$(odds ratio)) or depleted (negative $log_2$(odds ratio)) phyla.

| InDel Class | Phylum | Direction | Count | $log_2$(odds ratio) | p-value | FDR (p-value) |
|---|---|---|---|---|---|---|
| A | Actinomycetota | enriched | 2937 (86%) | 3.9 | 0 | 0 |
| | Methanobacteriota | enriched | 55 (1.6%) | 1.8 | 3.90E-15 | 1.20E-13 |
| | Bacteroidota | depleted | 0 (0%) | -9.7 | 1.90E-06 | 4.60E-05 |
| | Spirochaetota | depleted | 0 (0%) | -6.4 | 0.0018 | 0.024 |
| | Myxococcota | depleted | 1 (0.029%) | -4.7 | 6.80E-05 | 0.001 |
| | Bacillota | depleted | 108 (3.2%) | -3.7 | 4.80E-150 | 2.90E-148 |
| | Acidobacteriota | depleted | 3 (0.088%) | -3.4 | 8.80E-06 | 0.00015 |
| B | Chlamydiota | enriched | 15 (0.19%) | 2.2 | 4.80E-05 | 0.00058 |
| | Actinomycetota | enriched | 4148 (53%) | 1.5 | 0 | 0 |
| | Mycoplasmatota | enriched | 34 (0.44%) | 0.93 | 0.0018 | 0.02 |
| | Bacteroidota | enriched | 876 (11%) | 0.24 | 3.40E-05 | 0.00049 |
| | Methanobacteriota | depleted | 1 (0.013%) | -5.2 | 9.10E-06 | 0.00016 |
| | Spirochaetota | depleted | 10 (0.13%) | -3.3 | 1.50E-13 | 6.00E-12 |
| | Verrucomicrobiota | depleted | 14 (0.18%) | -1.9 | 1.30E-06 | 3.10E-05 |
| | Bacillota | depleted | 906 (12%) | -1.8 | 1.10E-243 | 6.90E-242 |
| | Planctomycetota | depleted | 12 (0.15%) | -1.7 | 3.60E-05 | 0.00049 |
| C | Fibrobacterota | enriched | 55 (0.53%) | 6.7 | 2.40E-08 | 2.50E-07 |
| | Candidatus Aminicenantota | enriched | 9 (0.087%) | 5.7 | 0.0065 | 0.047 |
| | Dictyoglomota | enriched | 8 (0.077%) | 3.2 | 0.002 | 0.016 |
| | Rhodothermota | enriched | 21 (0.2%) | 3 | 1.30E-06 | 1.20E-05 |
| | Cyanobacteriota | enriched | 76 (0.73%) | 2.9 | 7.90E-20 | 1.20E-18 |
| | Actinomycetota | depleted | 938 (9.1%) | -2.9 | 0 | 0 |
| | Spirochaetota | depleted | 24 (0.23%) | -2.6 | 5.00E-18 | 6.10E-17 |
| | Chloroflexota | depleted | 80 (0.77%) | -1.5 | 1.90E-18 | 2.60E-17 |
| | Acidobacteriota | depleted | 68 (0.66%) | -0.8 | 3.30E-05 | 0.00029 |
| | Bacillota | depleted | 2091 (20%) | -0.77 | 6.40E-83 | 1.90E-81 |
| D | Deinococcota | enriched | 16 (0.69%) | 3.1 | 3.50E-12 | 1.40E-10 |
| | Actinomycetota | enriched | 1760 (75%) | 2.8 | 0 | 0 |
| | Myxococcota | enriched | 54 (2.3%) | 1.3 | 6.00E-10 | 1.80E-08 |
| | Acidobacteriota | enriched | 49 (2.1%) | 1.2 | 9.30E-08 | 2.30E-06 |
| | Bacteroidota | depleted | 0 (0%) | -9.1 | 7.80E-06 | 0.00016 |
| | Bacillota | depleted | 3 (0.13%) | -8.1 | 8.50E-26 | 5.20E-24 |

| | | | | | | |
|---|---|---|---|---|---|---|
| **E** | Actinomycetota | enriched | 2342 (98%) | 7 | 2.70E-217 | 3.30E-215 |
| | Bacillota | depleted | 0 (0%) | -11 | 8.10E-08 | 3.30E-06 |
| | Bacteroidota | depleted | 0 (0%) | -9.2 | 7.20E-06 | 0.00018 |
| | Pseudomonadota | depleted | 33 (1.4%) | -4.1 | 2.30E-58 | 1.40E-56 |
| | Chloroflexota | depleted | 4 (0.17%) | -3.4 | 8.50E-07 | 2.60E-05 |
| | Myxococcota | depleted | 5 (0.21%) | -2.2 | 0.00029 | 0.0058 |
| **F** | candidate division WOR-3 | enriched | 16 (1.4%) | 10 | 1.00E-06 | 4.90E-06 |
| | Candidatus Daviesiibacteriota | enriched | 12 (1.1%) | 9.7 | 3.00E-06 | 1.40E-05 |
| | Candidatus Beckwithiibacteriota | enriched | 10 (0.89%) | 9.5 | 5.80E-06 | 2.50E-05 |
| | Candidatus Collieribacteriota | enriched | 28 (2.5%) | 8.6 | 1.60E-19 | 1.60E-18 |
| | Candidatus Fermentibacterota | enriched | 5 (0.45%) | 8.5 | 6.30E-05 | 0.00025 |
| | Bacillota | depleted | 3 (0.27%) | -7 | 1.70E-19 | 1.60E-18 |
| | Actinomycetota | depleted | 15 (1.3%) | -5.2 | 1.10E-45 | 2.10E-44 |
| | Spirochaetota | depleted | 0 (0%) | -4.7 | 0.022 | 0.05 |
| | Pseudomonadota | depleted | 21 (1.9%) | -3.5 | 2.50E-29 | 3.10E-28 |
| | Acidobacteriota | depleted | 2 (0.18%) | -2.2 | 0.015 | 0.035 |
| **G** | Candidatus Kaiseribacteriota | enriched | 51 (0.46%) | 8 | 9.70E-05 | 0.00069 |
| | Thermoproteota | enriched | 21 (0.19%) | 6.7 | 0.0011 | 0.0059 |
| | Candidatus Giovannoniibacteriota | enriched | 17 (0.15%) | 6.4 | 0.0019 | 0.0095 |
| | Synergistota | enriched | 14 (0.13%) | 6.2 | 0.003 | 0.014 |
| | Candidatus Yonathiibacteriota | enriched | 8 (0.072%) | 5.4 | 0.01 | 0.038 |
| | Fibrobacterota | depleted | 1 (0.009%) | -3.9 | 0.0011 | 0.0059 |
| | Fidelibacterota | depleted | 3 (0.027%) | -3.5 | 7.30E-06 | 5.90E-05 |
| | Actinomycetota | depleted | 834 (7.5%) | -3.3 | 0 | 0 |
| | Ignavibacteriota | depleted | 6 (0.054%) | -3 | 2.40E-07 | 2.00E-06 |
| | Methanobacteriota | depleted | 13 (0.12%) | -2.7 | 3.90E-11 | 4.70E-10 |

**Supplementary Table 16.** Statistical enrichment calculations of the broad biological categories of auxiliary domains associated with catalytic domain InDel classes. Enrichment for each class/category combination is relative to the global dataset.

| InDel Class | Broad category | Direction | count (% of total InDel Class) | $log_2$(odds ratio) | FDR (p value) |
|---|---|---|---|---|---|
| A | Carbohydrate binding / metabolism | enriched | 3395 (69.9%) | 2.2 | 0.0e+00 |
| | Transport / secretion system | enriched | 505 (10.4%) | 1.4 | 2.2e-72 |
| | Cell wall / peptidoglycan binding or remodelling | depleted | 1 (0.0%) | -9.0 | 3.720663e-14 |
| | Nucleic acid / protein binding | depleted | 0 (0.0%) | -4.3 | 4.8e-02 |
| | Enzymatic / catalytic function | depleted | 43 (0.9%) | -2.4 | 3.7e-27 |
| B | Stress response / defence | enriched | 12 (0.1%) | 5.0 | 1.1e-04 |
| | Structural / scaffolding | enriched | 2613 (24.2%) | 1.1 | 3.6e-164 |
| | Transport / secretion system | enriched | 760 (7.0%) | 0.74 | 6.6e-30 |
| | Cell wall / peptidoglycan binding or remodelling | depleted | 12 (0.1%) | -7.3 | 3.2e-71 |
| | Nucleic acid / protein binding | depleted | 0 (0.0%) | -5.6 | 7.8e-03 |
| | Signal transduction / regulatory | depleted | 1 (0.0%) | -3.5 | 4.3e-03 |
| C | Signal transduction / regulatory | enriched | 63 (0.4%) | 5.3 | 2.1e-11 |
| | Nucleic acid / protein binding | enriched | 85 (0.5%) | 4.7 | 3.2e-17 |
| | DUF/unknown | enriched | 277 (1.7%) | 1.1 | 1.4e-18 |
| | Cell wall / peptidoglycan binding or remodelling | depleted | 181 (1.1%) | -4.2 | 0.0e+00 |
| | Enzymatic / catalytic function | depleted | 241 (1.5%) | -1.9 | 9.1e-83 |
| | Transport / secretion system | depleted | 610 (3.8%) | -0.52 | 3.1e-14 |
| D | Carbohydrate binding / metabolism | enriched | 1695 (57.1%) | 1.2 | 2.0e-109 |
| | Structural / scaffolding | enriched | 704 (23.7%) | 0.81 | 2.1e-35 |
| | Transport / secretion system | enriched | 212 (7.1%) | 0.63 | 1.1e-08 |
| | Cell wall / peptidoglycan binding or remodelling | depleted | 1 (0.0%) | -8.2 | 7.2e-12 |
| | Enzymatic / catalytic function | depleted | 9 (0.3%) | -3.9 | 5.6e-16 |
| | DUF/unknown | depleted | 6 (0.2%) | -2.4 | 3.9e-05 |
| E | Structural / scaffolding | enriched | 1029 (37.2%) | 1.8 | 8.0e-204 |
| | Carbohydrate binding / metabolism | enriched | 1475 (53.3%) | 0.99 | 9.3e-68 |
| | Transport / secretion system | enriched | 168 (6.1%) | 0.35 | 4.5e-03 |
| | Cell wall / peptidoglycan binding or remodelling | depleted | 0 (0.0%) | -9.7 | 5.0e-06 |
| | Enzymatic / catalytic function | depleted | 0 (0.0%) | -8.0 | 1.8e-04 |
| | DUF/unknown | depleted | 0 (0.0%) | -6.0 | 4.45e-03 |
| F | Metal / cofactor binding | enriched | 1 (0.1%) | 5.7 | 1.1e-03 |
| | Transport / secretion system | enriched | 239 (24.2%) | 2.8 | 8.7e-134 |
| | DUF/unknown | enriched | 39 (3.9%) | 2.0 | 1.3e-15 |
| | Carbohydrate binding / metabolism | depleted | 36 (3.6%) | -4.0 | 8.3e-60 |
| | Cell wall / peptidoglycan binding or remodelling | depleted | 32 (3.2%) | -2.1 | 1.3e-15 |
| G | Cell wall / peptidoglycan binding or remodelling | enriched | 6208 (47.2%) | 7.2 | 0.0e+00 |
| | Enzymatic / catalytic function | enriched | 1323 (10.0%) | 2.3 | 8.0e-269 |
| | Other / unclassified | enriched | 3394 (25.8%) | 1.1 | 2.8e-194 |
| | Signal transduction / regulatory | depleted | 0 (0.0%) | -5.5 | 8.4e-03 |
| | Transport / secretion system | depleted | 31 (0.2%) | -4.9 | 4. 9e-78 |
| | Structural / scaffolding | depleted | 171 (1.3%) | -4.3 | 0.0e+00 |

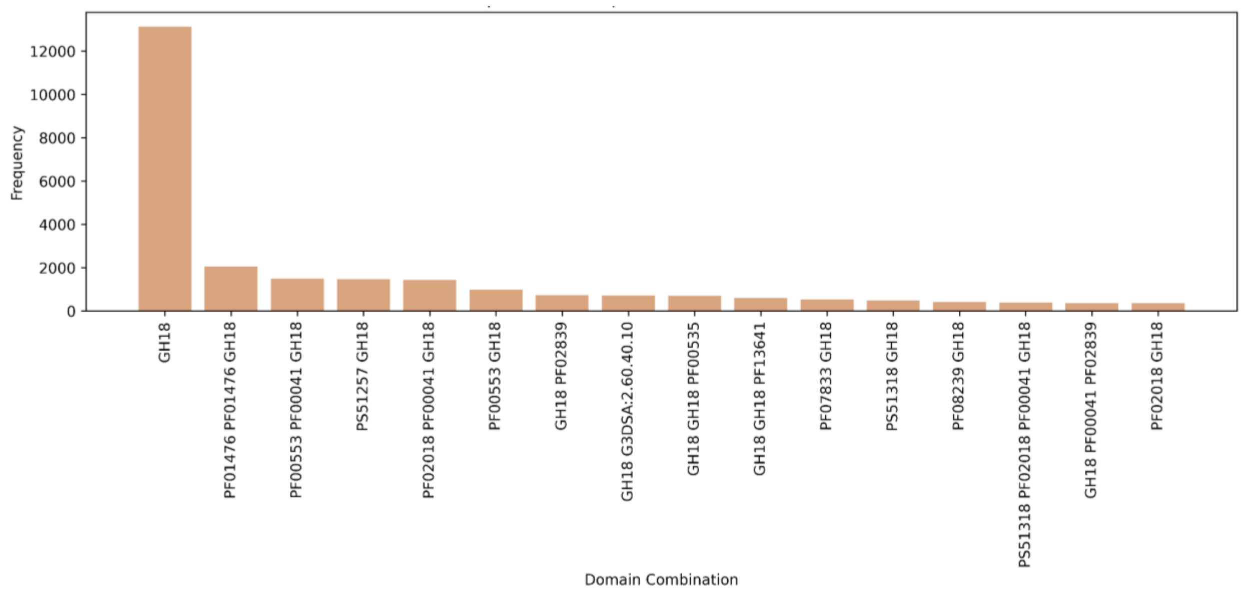


**Supplementary Figure 1.** Frequency of GH18 only sequences and the top 15 most common multidomain architectures observed across the dataset.

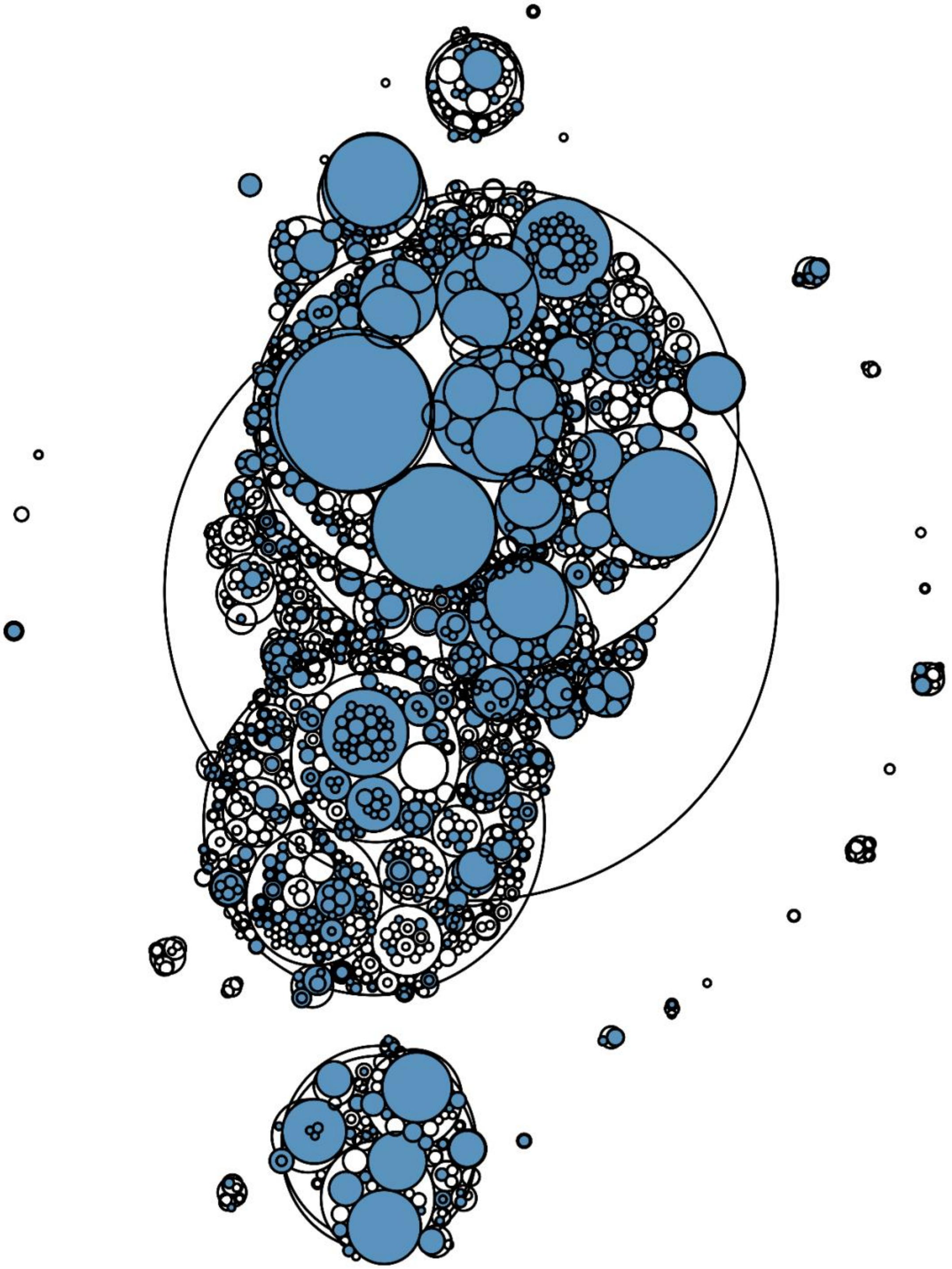

**Supplementary Figure 2.** The SSN depicted in Figure 1 of the main text with clusters at the highest bitscore threshold coloured blue to indicate the presence of a sequence comprising only a single catalytic domain.

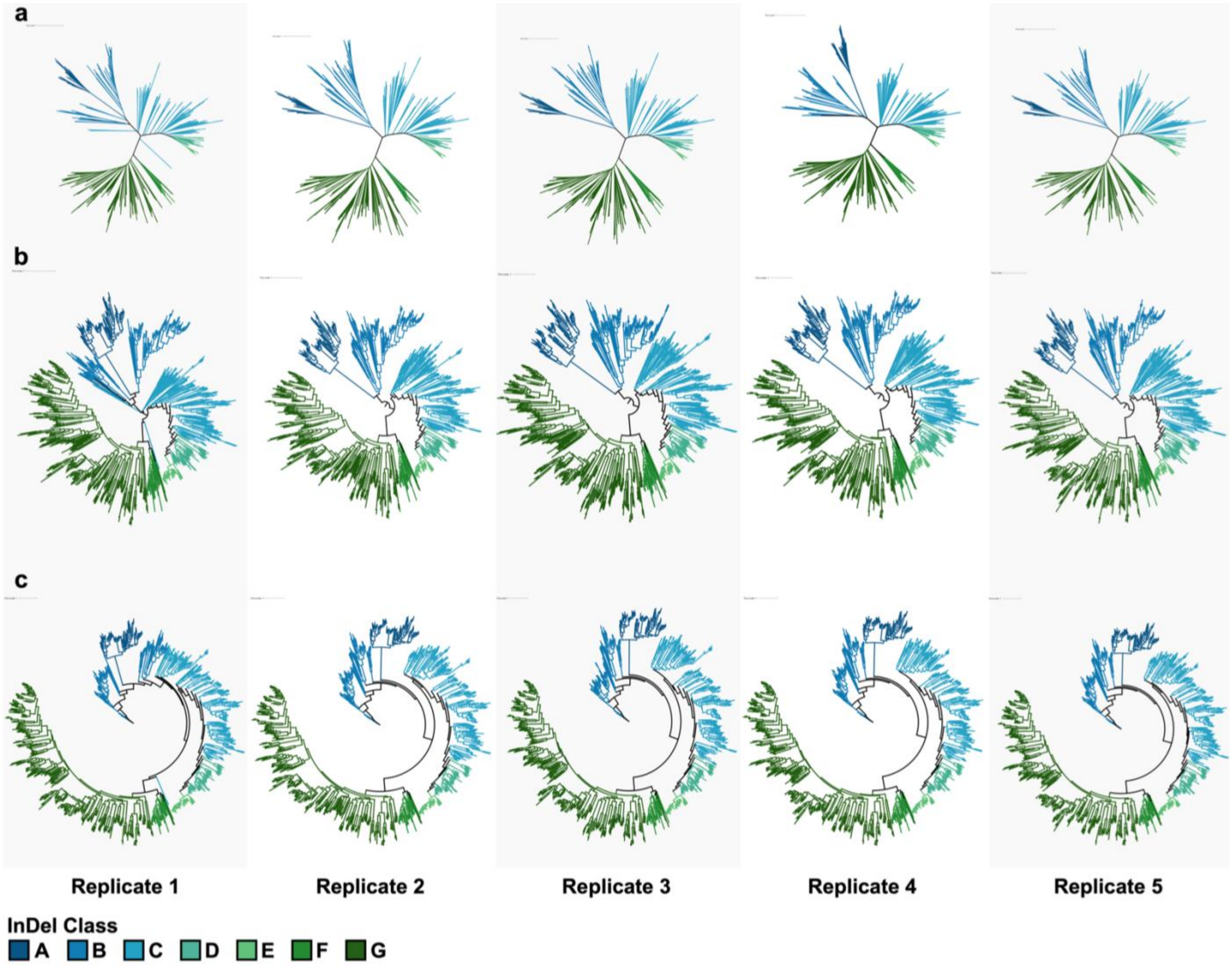


**Supplementary Figure 3.** Replicate phylogenies independently inferred from the same multiple sequence alignment of GH18 catalytic domains using IQTREE3. Trees are shown either in (a) an unrooted orientation, (b) midpoint rooted or (c) rooted using the minimal ancestral deviation method. All trees are coloured with respect to the InDel Classes discussed in the main text and annotated in Figure 3a.

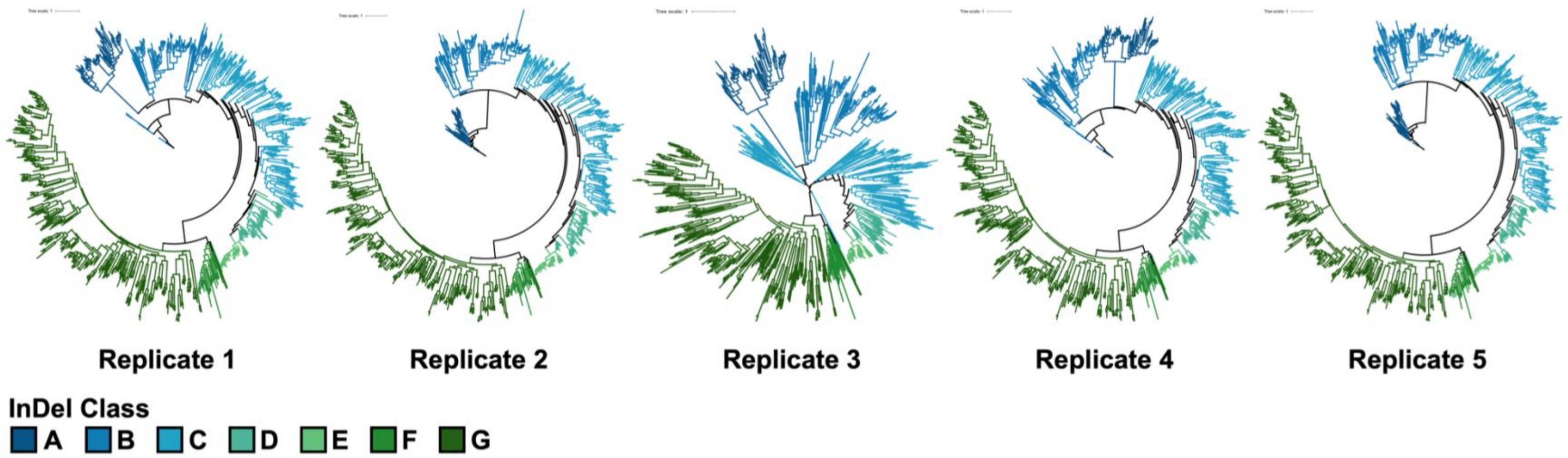


**Supplementary Figure 4.** Replicate phylogenies independently inferred from the same multiple sequence alignment of GH18 catalytic domains using IQTREE3 under a non-reversible substitution model. Of these, ModelFinder (inbuilt in the IQTREE3 pipeline) selected nq.PFAM+F+R10 as the model of best fit independently across all replicates. All trees are coloured with respect to the InDel Classes discussed in the main text and annotated in Figure 3a.

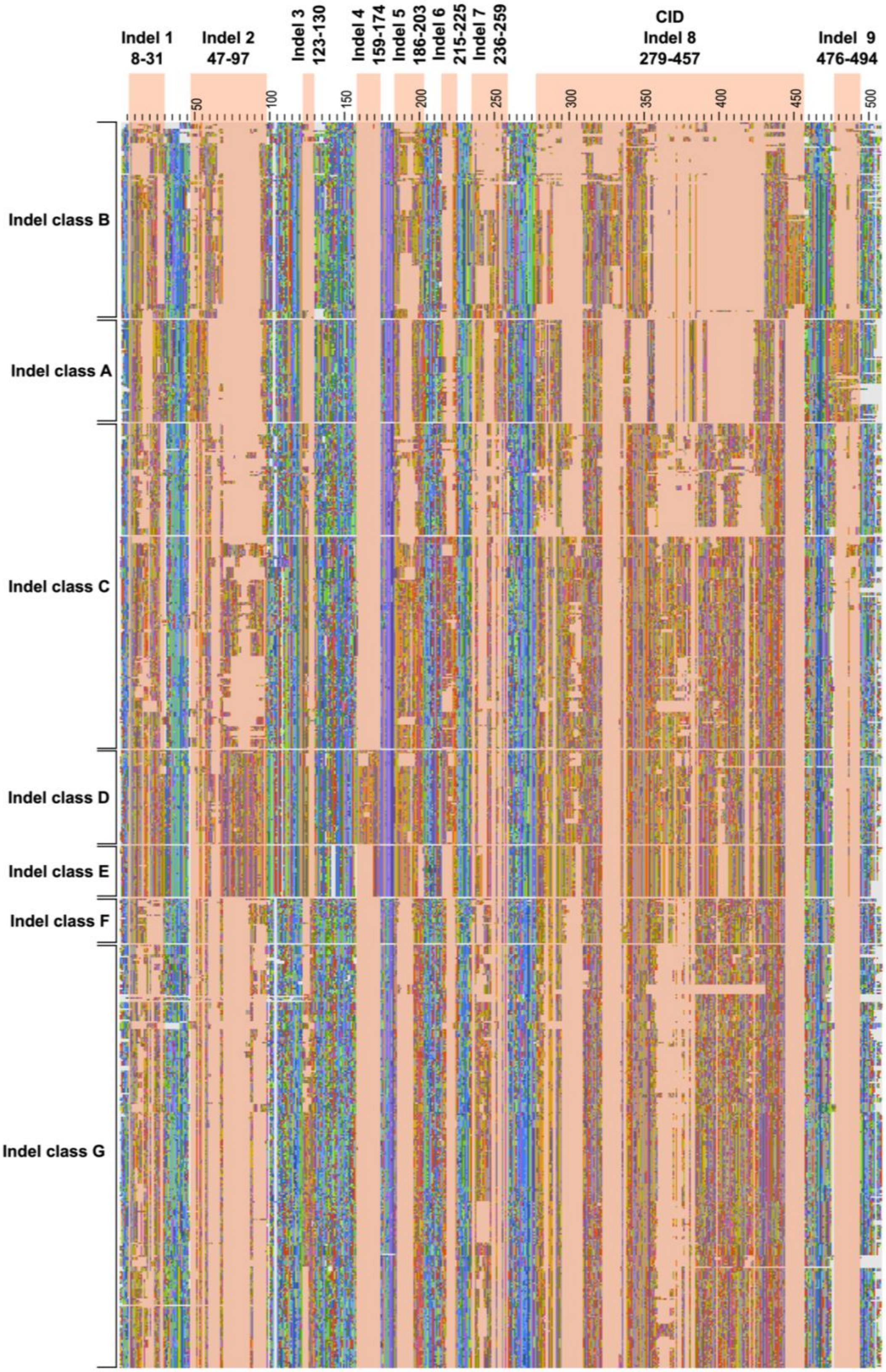


**Supplementary Figure 5.** Annotated multiple sequence alignment from which the phylogeny displayed in Figure 3a in the main text with each Indel Class indicated and the insertion/deletion positions that define them.

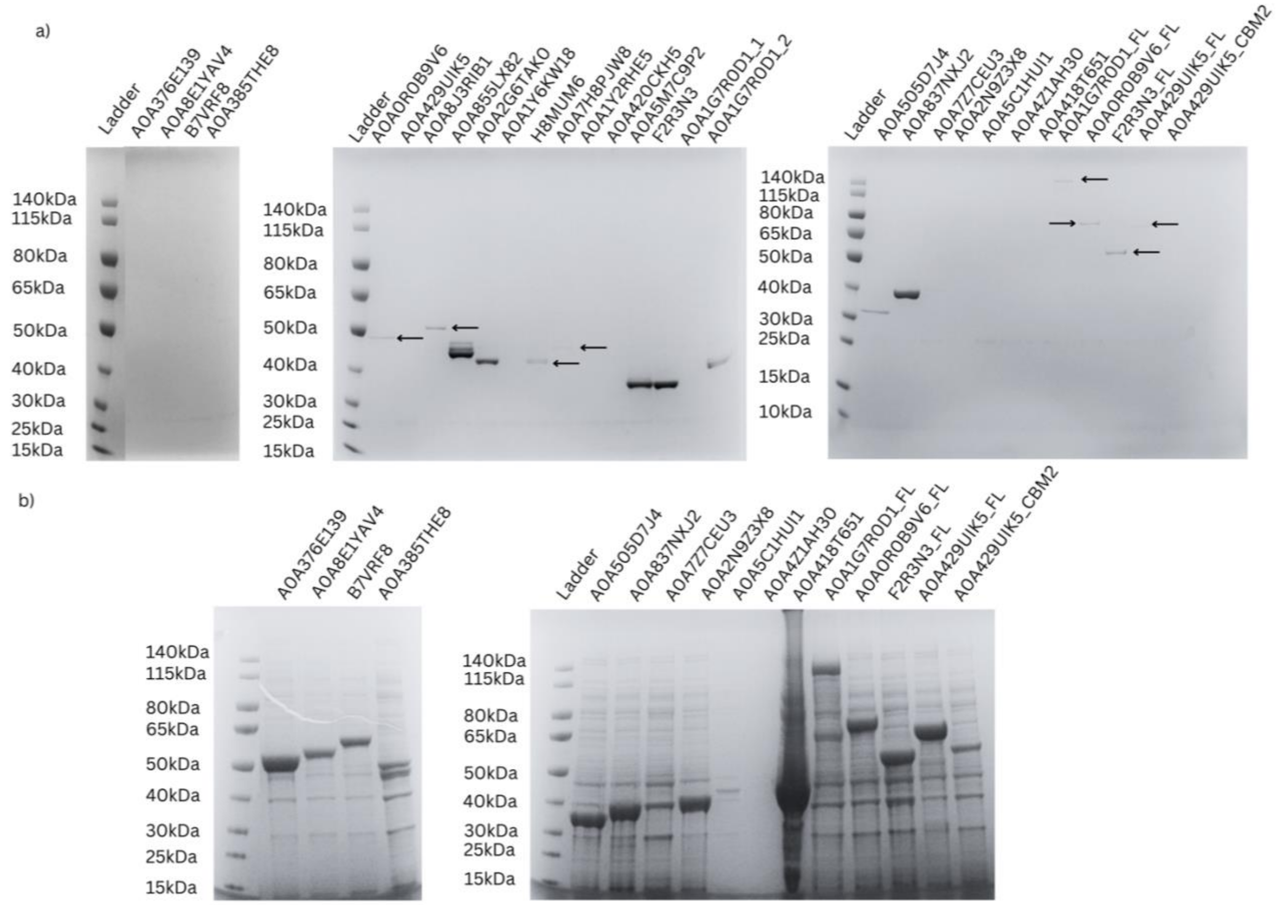


**Supplementary Figure 6. High-throughput expression and solubility screening of heterologously expressed chitinases. (a)** SDS–PAGE analysis of chitinases expressed in 96-well format and purified by $Ni^{2+}$–NTA affinity chromatography; in each case, the predominant band corresponded to the predicted molecular weight (indicated by arrows). **(b)** Representative SDS–PAGE showing the insoluble fractions of a subset of chitinases, illustrating that all constructs were detected in the insoluble fraction, irrespective of whether correctly folded soluble protein was recovered; purified proteins again migrated at their predicted molecular weights.

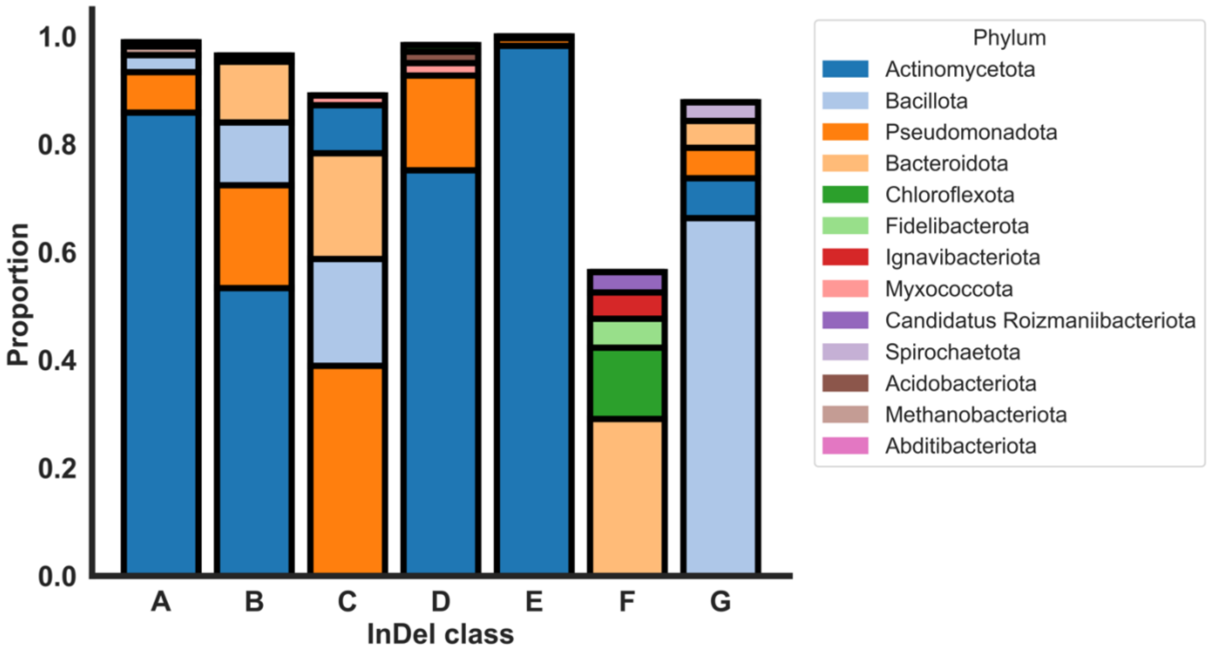


**Supplementary Figure 7.** Phylum-level composition of sequences across catalytic InDel classes. Stacked bar plots show, for each catalytic InDel class (A–G), the proportional representation of the top five phyla contributing sequences to that class. Colours are consistent across InDel classes and are assigned to phyla as per the legend in figure.

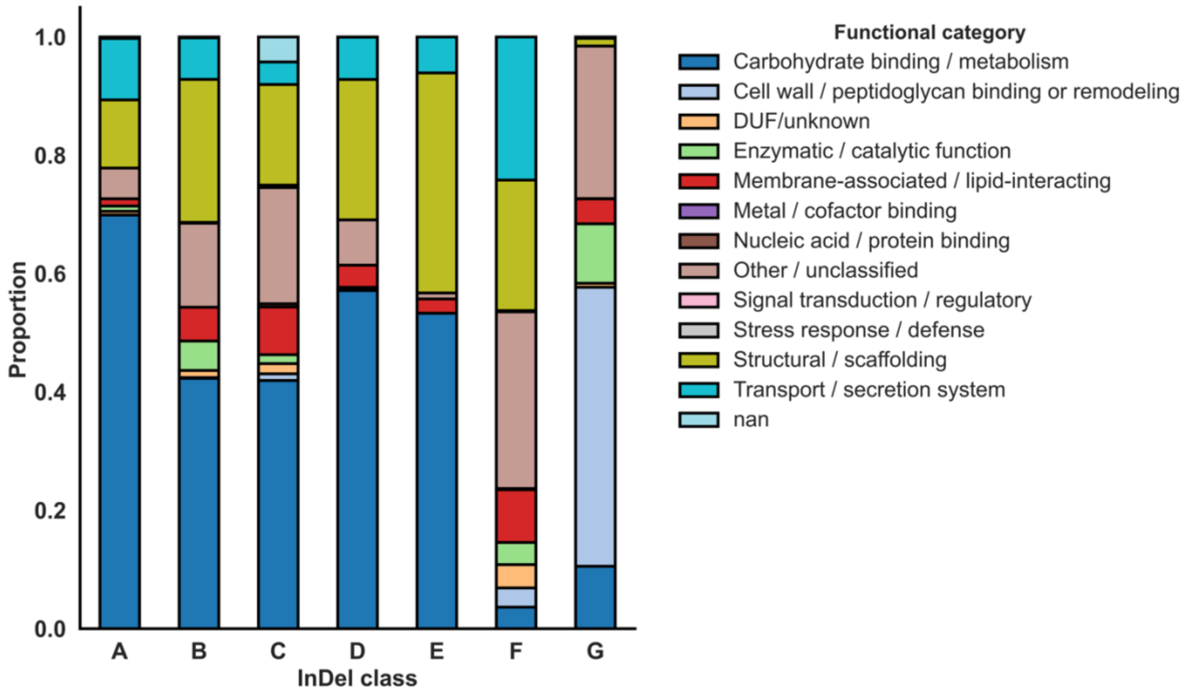


**Supplementary Figure 8.** Stacked bar plots show, for each catalytic InDel class (A–G), the proportion of associated auxiliary domains assigned to each broad functional category (see legend in figure). Bar segments within each InDel class sum to 1, illustrating relative rather than absolute contributions of each functional category; auxiliary domains lacking a functional category annotation are omitted for clarity.

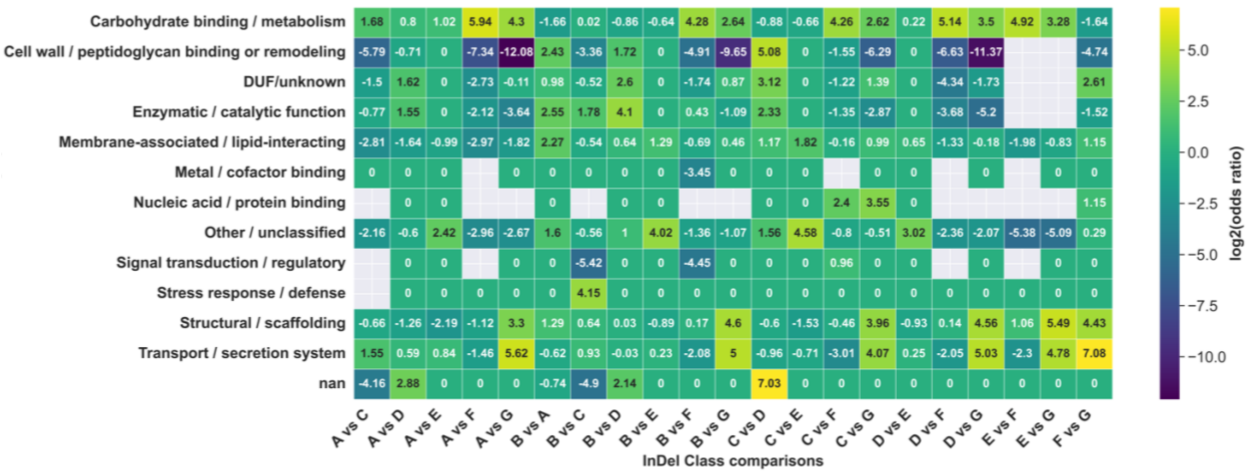


**Supplementary Figure 9.** Heatmap of $log_2$(odds ratio) pairwise comparisons of all InDel classes and auxiliary domain biological categories.
